# PDZD8 promotes autophagy at ER-Lysosome contact sites to regulate synaptogenesis

**DOI:** 10.1101/2023.10.30.564828

**Authors:** Rajan S. Thakur, Kate M. O’Connor-Giles

## Abstract

Building synaptic connections, which are often far from the soma, requires coordinating a host of cellular activities from transcription to protein turnover, placing a high demand on intracellular communication. Membrane contact sites (MCSs) formed between cellular organelles have emerged as key signaling hubs for coordinating an array of cellular activities. We have found that the endoplasmic reticulum (ER) MCS tethering protein PDZD8 is required for activity-dependent synaptogenesis. PDZD8 is sufficient to drive ectopic synaptic bouton formation through an autophagy-dependent mechanism and required for basal synapse formation when autophagy biogenesis is limited. PDZD8 functions at ER-late endosome/lysosome (LEL) MCSs to promote lysosome maturation and accelerate autophagic flux. Mutational analysis of PDZD8’s SMP domain further suggests a role for lipid transfer at ER-LEL MCSs. We propose that PDZD8-dependent lipid transfer from ER to LELs promotes lysosome maturation to increase autophagic flux during periods of high demand, including activity-dependent synapse formation.

**GRAPHICAL ABSTRACT:** 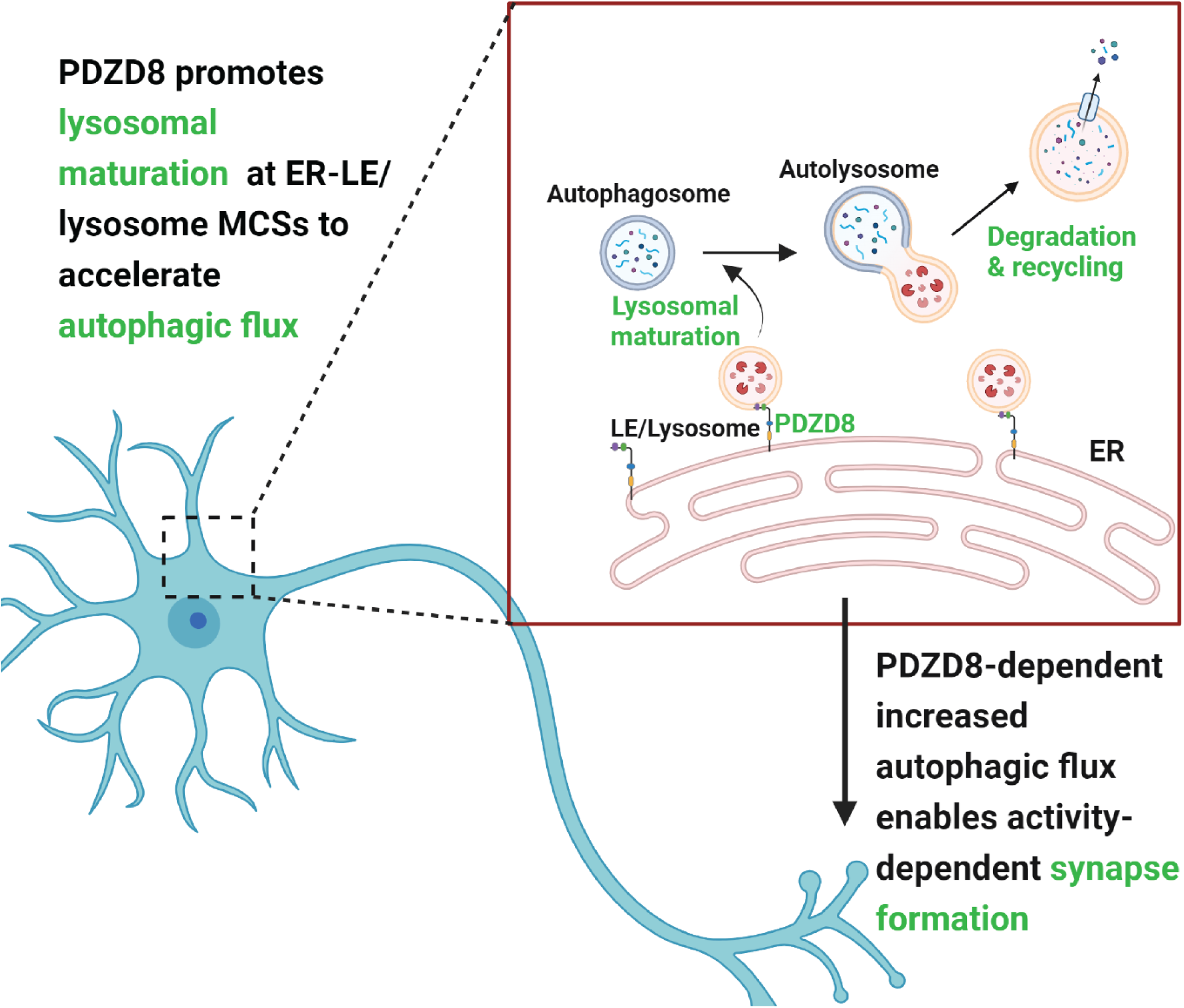

## INTRODUCTION

Neurons are large, polarized cells that must coordinate the formation and long-term function of synaptic connections. The great distances between cell bodies and synapses pose a unique challenge for intracellular communication in neurons. Over the last decade, membrane contact sites (MCSs), sites of close apposition between cellular organelles, have been increasingly recognized as key signaling hubs for coordinating a vast array of cellular activities. MCSs are formed by tethering proteins residing in organelle membranes^1–5^.

In an ongoing screen for conserved regulators of synaptic growth, we identified the resident ER tethering protein PDZD8 as a positive regulator of activity-dependent synapse formation. PDZD8 is an SMP (synaptotagmin-like mitochondrial-lipid-binding) domain protein belonging to the tubular lipid-binding protein (TULIP) superfamily of lipid transfer proteins. The ER is the largest membrane bound organelle, forming a network that extends throughout the entire cell, including the distalmost compartments of neurons^3^. The ER interacts with virtually all other cellular organelles through MCSs ^1,6–8^, enabling the non-vesicular exchange of lipids, ions, and other metabolites, and function as communication sites for regulating diverse cellular functions^2–5,9,10^.

PDZD8 has come to recent attention for its role in tethering multiple ER MCSs^11–16^. PDZD8 was first reported to regulate dendritic Ca^2+^ dynamics in cultured cortical neurons by promoting ER-mitochondria tethering ^16^. More recent studies indicate that a primary PDZD8 function is tethering ER–late endosome/lysosome (LEL) MCSs through interactions with Rab7 and Protrudin^12–15^. In mouse primary cortical neurons and PC12 cells, PDZD8 promotes axon maintenance and neurite outgrowth, respectively^12,13^, and a recent study in *Drosophila* observed beneficial effects of RNAi knockdown of PDZD8 in aging^11^. Consistent with critical roles in the nervous system, two nonsense variants of *PDZD8* were recently identified as the cause of autosomal recessive syndromic intellectual disability in multiple families^17^.

Here, we investigate the *in vivo* role of PDZD8 in nervous system development using *Drosophila* as a model. We find that PDZD8 is broadly expressed in neurons, where it predominantly localizes to ER-LEL MCSs. PDZD8 is required for activity-induced synaptogenesis and sufficient to promote ectopic synaptic bouton formation. We find that PDZD8 promotes synatogenesis through a macroautophagy-dependent mechanism. Macroautophagy (hereafter referred to as autophagy) is a conserved pathway culminating in the degradation of damaged organelles, protein aggregates, and long-lived proteins in autolysosomes^18^. Our studies suggest that PDZD8 accelerates autophagic flux by promoting lysosome maturation and, thus, the degradative function of autolysosomes. Disruption of PDZD8 lipid-binding disrupts PDZD8 co-localization with LELs, its role in promoting autolysosome turnover, and its ability to induce ectopic synapse formation. Mutation of key residues in the SMP lipid-transfer domain also diminish PDZD8-induced ectopic synapse formation, implicating PDZD8 lipid binding and transfer activity at ER-LEL MCSs. Together our findings support the model that PDZD8 increases autophagic flux to promote synaptic growth during periods of high demand such as activity-dependent synapse formation.

## RESULTS

### PDZD8 is required for activity-dependent synaptogenesis

In an ongoing screening effort for new synaptic genes, we identified *PDZD8* as a candidate based on its transcriptional spatio-temporal expression profiles, which specifically corresponds to periods of peak synaptogenesis in the developing *Drosophila* nervous system ^19^. To investigate PDZD8’s potential role in nervous system development, we used CRISPR to generate null and endogenously tagged alleles (see Materials and methods) and quantified baseline and activity-dependent synaptic bouton formation at the well-characterized glutamatergic larval neuromuscular junction (NMJ). At NMJs, a single motor neuron generally innervates a single muscle cell and forms a stereotyped number of synaptic boutons, each containing ∼10 individual active zones aligned with postsynaptic glutamate receptor clusters^20^. Synaptic bouton formation occurs normally in the absence of PDZD8 at the standard rearing temperature of 25℃, indicating that PDZD8 is dispensable for basal synapse formation under these conditions (Fig. 1A,B). We next investigated activity-dependent synaptogenesis. Previous studies have shown that rearing larvae at elevated temperature leads to activity-induced NMJ expansion due to increased larval locomotion and synaptic activity^21 22^. In wild-type controls, we see an ∼40% increase in the number of synaptic boutons in animals reared at 31℃ vs. 25℃ (Fig. 1A,B). In contrast, no activity-dependent bouton formation is observed in *PDZD8^KO^* (Fig. 1A,B). To confirm that loss of *PDZD8* is responsible for the absence of activity-dependent synaptic bouton formation, we restored a single copy of *PDZD8* under endogenous regulation using a genomic duplication ^23^. Restoring PDZD8 function raises bouton number to control levels at 31℃ (Fig. 1C,D), confirming a requirement in activity-dependent synapse formation *in vivo*.

**Figure 1.**
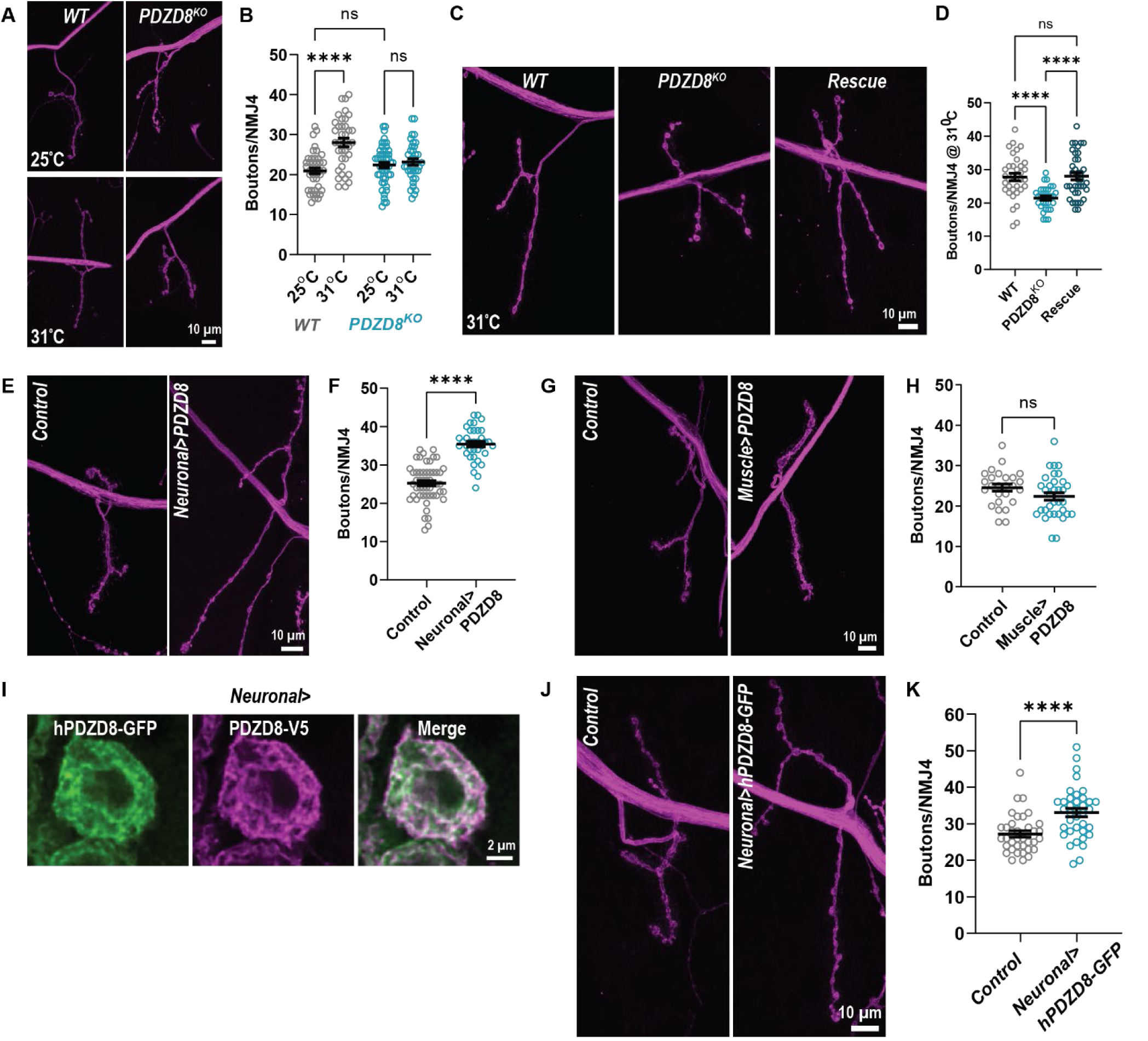
PDZD8 is required for activity-dependent synaptogenesis. **(A-D)** PDZD8 is dispensable for basal synaptic growth (25^0^C) but required for activity induced synaptic growth (31^0^C). Representative confocal images of NMJ labeled with FITC-conjugated anti-HRP (magenta **(A,C)** and bouton quantification **(B,D)** in the indicated genotypes. Scale bar=10 μm. **(E-H)** Neuronal overexpression of *PDZD8* induces ectopic synaptic growth, whereas muscle overexpression has no effect. Representative confocal images of NMJs labeled with FITC-conjugated anti-HRP (magenta) **(E,G)** and bouton quantification **(F,H)** in the indicated genotypes. *elav-Gal4* was used to drive expression in the neurons and *24B-Gal4* is used to drive expression in muscle. Scale bar=10 μm. **(I)** hPDZD8 expression in neurons completely overlaps with PDZD8. SoRa spinning disk deconvolved Z-projections of a *elav>PDZD8-V5, hPDZD8-GFP* larval neuronal cell body co-labeled with antibodies against GFP (green) and V5 (magenta). Scale bar=2 μm. **(J-K)** *hPDZD8* misexpression in neurons leads to ectopic synaptic growth. Representative confocal images of NMJs labeled with FITC-conjugated anti-HRP (magenta) **(J)** and bouton quantification **(K)** in the indicated genotypes. Scale bar=10 μm. Data is represented as mean ± SEM with significance calculated by ANOVA followed by Tukey’s multiple comparisons test **(B,D)**, Unpaired t test **(F,H)**, or Mann-Whitney U test **(K)**. ****p < 0.0001, n.s. = not significant.

We next sought to determine if PDZD8 is sufficient to promote synaptogenesis by driving ectopic *PDZD8* expression in neurons under the control of the pan-neuronal driver *elav-Gal4*. We observe significant ectopic bouton formation upon neuronal *PDZD8* overexpression, demonstrating that PDZD8 is sufficient to promote synaptogenesis (Fig. 1E,F). We further find that C-terminal tagging of PDZD8 does not affect overall protein function as pan-neuronal expression of *PDZD8-V5* also promotes ectopic bouton formation (Fig. S1A-C). In contrast, expression of PDZD8 in postsynaptic muscle under the control of the driver *24B-Gal4* has no effect on bouton number (Fig. 1G,H), indicating a presynaptic-specific role for PDZD8 in promoting synaptogenesis. To assess the conservation of PDZD8’s role in synaptogenesis, we expressed GFP-tagged human *PDZD8* (*hPDZD8-GFP*) in neurons. In neuronal cell bodies, hPDZD8-GFP fully overlaps with PDZD8-V5 in a reticular ER pattern (Fig. 1I), consistent with shared regulation, and similarly induces ectopic bouton formation (Fig. 1J,K). Together, our findings demonstrate a conserved presynaptic role for PDZD8 in promoting synaptogenesis.

### PDZD8 promotes synaptogenesis via autophagy

The morphology of expanded NMJs induced by *PDZD8* overexpression in neurons, with long branches of small boutons, is strikingly similar to the synaptic overgrowth observed when autophagy is increased ^24^, prompting us to investigate potential links between PDZD8 and autophagy. In neurons, autophagy is a constitutive process with diverse roles in synapse formation and maintenance across species^18,24–33^. Neuronal activity also induces autophagy at synapses to maintain protein quality control^34^. Based on the morphological similarity between NMJs overexpressing *PDZD8* or *atg1*, a critical autophagy biogenesis gene^35,36^, we hypothesized that the ectopic bouton formation we observed upon *PDZD8* overexpression may be due to increased autophagy. To test this, we reduced the levels of *atg1* while overexpressing *PDZD8*. We found that loss of a single copy of *atg1* in an otherwise wild-type background does not impact bouton formation (Fig. 2A,B). However, loss of single copy of *atg1* fully suppresses *PDZD8*-induced bouton formation (Fig. 2A,B), indicating that PDZD8-induced synaptic growth requires autophagy.

**Figure 2.**
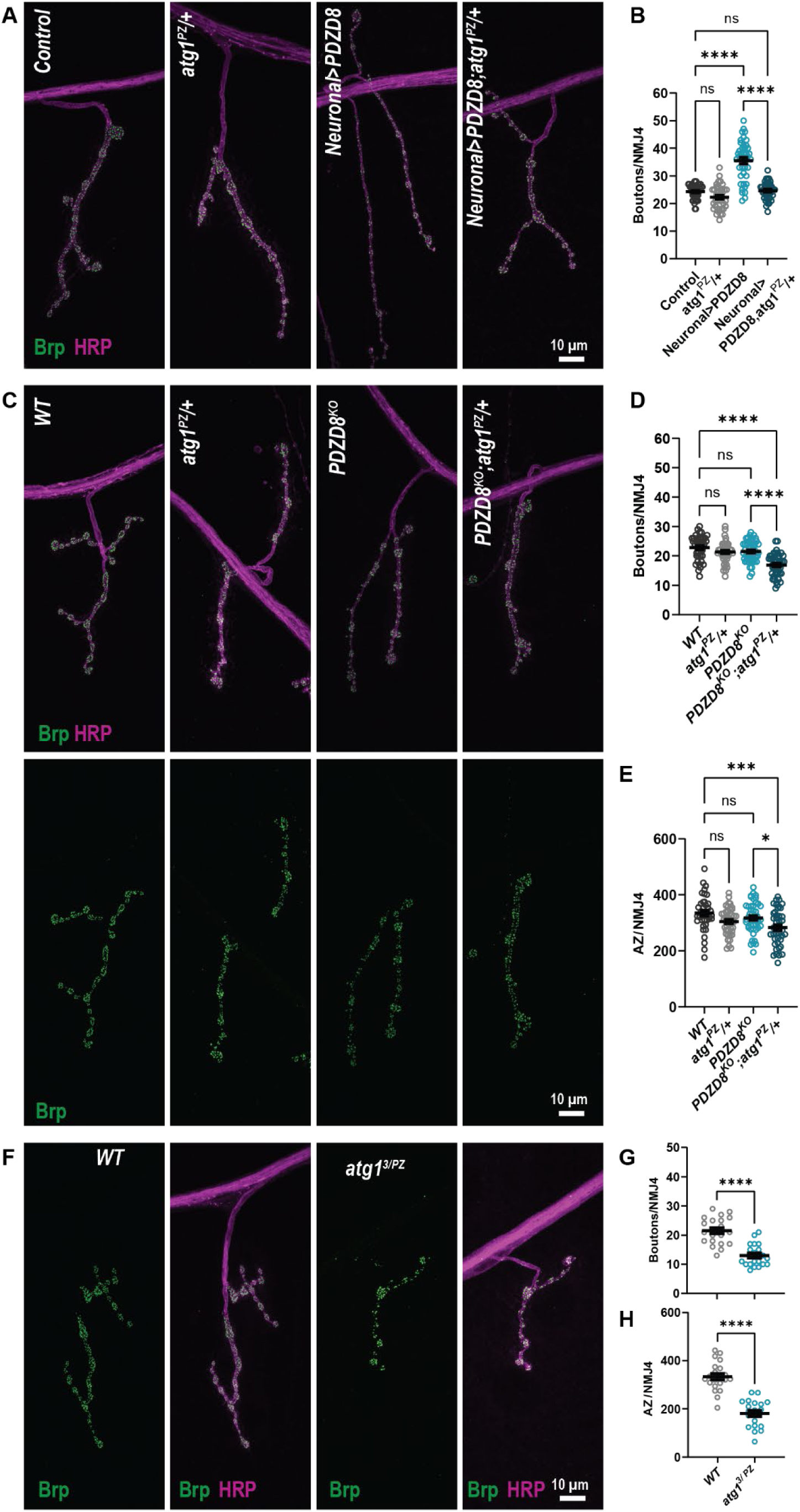
PDZD8 promotes synaptogenesis via autophagy. **(A-D)** Downregulation of autophagy completely suppresses PDZD8-induced ectopic synaptic growth and dominantly enhances the loss of *PDZD8,* leading to basal synaptic undergrowth. Representative confocal images of NMJs co-labeled with antibodies against active zone marker Brp (green) and FITC-conjugated anti-HRP (magenta) **(A,C)** and bouton quantification **(B,D)** in the indicated genotypes. *elav-Gal4* was used to drive *PDZD8* expression in neurons. **(C,E)** PDZD8-mediated autophagy regulates synapse formation. The total number of AZs per NMJ4 in the indicated genotypes. **(F-H)** Autophagy promotes bouton and synapse formation. Representative confocal images **(F)** co-labeled with antibodies against active zone marker Brp (green) and FITC-conjugated anti-HRP (magenta) and bouton number per NMJ 4 **(G)** and the total number of AZs per NMJ4 **(H)** in the indicated genotypes. Scale bar=10 μm. Data is represented as mean ± SEM with significance calculated by Kruskal-Wallis followed by Dunn’s multiple comparisons test **(B)**, ANOVA followed by Tukey’s multiple comparisons test **(D,E)** or Unpaired t-test **(G,H)**. *p < 0.05,**p < 0.01, ***p < 0.001,****p < 0.0001, n.s. = not significant.

This observation motivated us to test if, in a sensitized background such as *atg1^PZ^/+* where levels of a key regulator of autophagy are limiting, PDZD8 is required for synaptogenesis. Indeed, reducing *atg1* copy number dominantly enhances loss of *PDZD8*, revealing reduced bouton number when autophagy is limited (Fig. 2C,D). We observe similar interactions with *syntaxin 17*, which encodes a SNARE protein required for autophagosome-lysosome fusion^37^ (Fig. S2A-D). These findings demonstrate that PDZD8 promotes autophagy to positively regulate synaptogenesis.

In addition to its role in promoting synaptic bouton formation at the NMJ^24 18^, autophagy also plays key roles in the formation and refinement of individual synapses across species, including in the mouse cortex, fly visual system, and worm NMJs ^28–32^. To investigate the role of PDZD8 and autophagy in the formation of individual synapses, we labeled presynaptic terminals with the active zone marker Brp. Similar to boutons, loss of *PDZD8* alone does not impact the number of active zones per NMJ (Fig. 2C,E). However, the number of active zones per NMJ is significantly decreased in *PDZD8^KO^; atg1^PZ^/+* (Fig. 2C,E), indicating that PDZD8 accelerates autophagy to promote synapse formation as well as bouton formation. This suggests a requirement for autophagy in active zone formation. However, while autophagy has previously been shown to promote bouton formation^24^, its role in individual motor synapse formation has not been investigated. As predicted, we observe significantly reduced active zone number in homozygous *atg1* mutants, demonstrating that autophagy promotes bouton and synapse formation at the NMJ (Fig. 2F-H). We next investigated whether PDZD8-mediated autophagy is sufficient to induce active zone formation and found that, in contrast to bouton formation, overexpression of *PDZD8* does not affect the total number of active zones per NMJ (data not shown). Together, these findings demonstrate key roles for PDZD8 and autophagy in the formation of boutons and individual motor synapses and raise the question of how PDZD8 promotes autophagy.

### PDZD8 is enriched in neurons and localizes to ER-LEL MCSs

To better understand the role of PDZD8 in promoting synaptogenesis, we next investigated PDZD8 cellular and subcellular localization in the nervous system. We generated an endogenously tagged allele of *PDZD8* with V5-mCherry incorporated at the C-terminus and confirmed expression of the correct size protein by western blot (Fig. S3A). In the larval nervous system, we observed PDZD8^V5-mCherry^ expression in cell bodies and the synaptic neuropil of the ventral ganglion (Fig. 3A). To determine which cell types express PDZD8, we co-labeled with antibodies against the neuronal marker Elav and glial marker Repo. We found that PDZD8 is expressed in all neuronal cell bodies, but not detectable above background in Repo+ glial cells, indicating that PDZD8 is primarily neuronal (Fig. 3B,C). PDZD8 is expressed at the larval NMJ, where it is mostly localized as discrete punctae throughout the presynaptic terminal (Fig. 3D).

**Figure 3.**
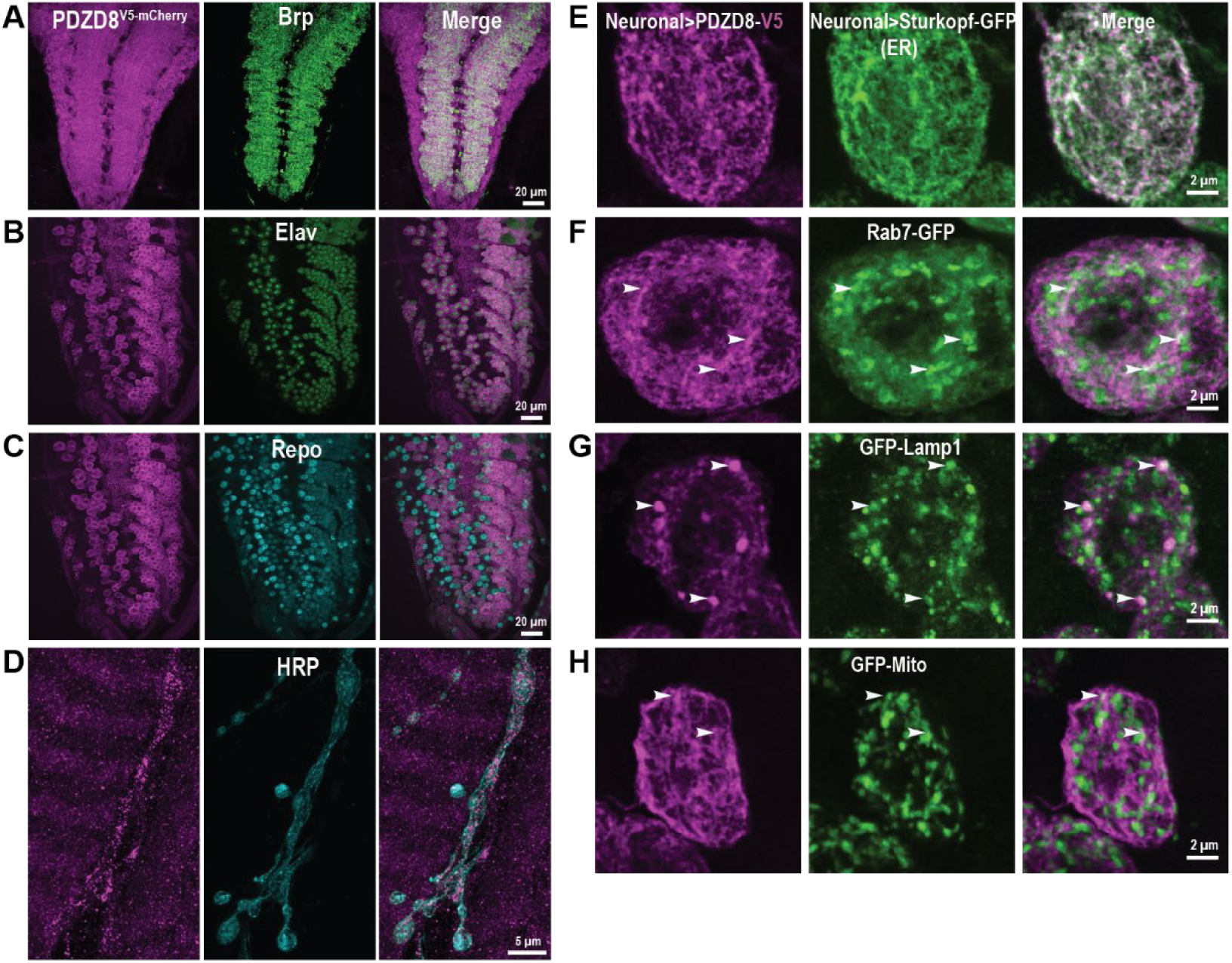
PDZD8 is enriched in neurons and localizes to ER-LEL MCSs. **(A-C)** PDZD8 is expressed in neurons and localized in cell bodies and at the synaptic neuropil. Confocal Z-projections of a *PDZD8^V^*^5^*^-mCherry^* larval ventral ganglia co-labeled with antibodies against mCherry (magenta) and the synaptic marker Brp in green **(A)**, neuronal cell marker Elav in green **(B)** and glial marker Repo in cyan **(C)**. Scale bar=20 μm. **(D)** PDZD8 is expressed at the NMJ. Confocal Z-projections of a *PDZD8^V^*^5^*^-mCherry^* larval NMJ 6/7 co-labeled with antibodies against mCherry (magenta) and the neuronal membrane marker HRP (cyan). Scale bar=5 μm. **(E-H)** PDZD8 exhibits a reticular organization in neurons, colocalizing with ER and multiple MCSs. SoRa spinning disk deconvolved single plane of an *elav-Gal4;PDZD8-V5* larval neuronal cell bodies co-labeled with antibodies against V5 (magenta) and neuronally expressed Sturkopf-GFP in green to label ER **(E)**, Rab7-GFP in green to label late endosomes **(F)**, GFP-Lamp1 in green to label lysosomes **(G)**, and GFP-Mito in green to label mitochondria **(H)**. White arrowheads highlight sites of colocalization in panels **F,G, and H**. Elav-Gal4 is used to drive endomembrane expression in the neurons. Scale bar=2 μm.

To better understand PDZD8 subcellular localization, we turned to super-resolution imaging via optical pixel reassignment. We found that both endogenously tagged PDZD8 and neuronally expressed PDZD8-V5 overlap extensively with the ER marker Sturkopf-GFP^38^. PDZD8^V5-mCherry^ exhibits a punctate pattern expected for ER proteins in fixed cells and similar to that observed in mammalian non-neuronal cells ^15,39^ (Fig. S3B), whereas neuronally expressed PDZD8-V5 exhibits a more reticular pattern (Fig. 3E and see Fig. 1I). Since PDZD8 encodes a protein reported to localize to multiple ER MCSs, we investigated PDZD8 co-localization with cellular organelles using LEL markers Rab7-GFP and GFP-Lamp1 and mitochondrial marker GFP-Mito (Fig. 3F-H and Fig. S3C-E). We observe significant colocalization of PDZD8 and LEL membranes consistent with recent studies indicating that PDZD8 is enriched at ER-LEL MCSs in non-neuronal cells ^12–15^. Notably, co-expression of PDZD8-V5 and GFP-Lamp1 leads to an increase in large lysosomal structures, similar to observations in mammalian cells^14,15^ (Fig. 3G). We further confirmed ER-lysosome MCS localization of PDZD8 by colabelling ER (RFP-KDEL) and lysosomes (GFP-Lamp1) along with PDZD8 (Fig. S4A,B). Consistent with prior observations of PDZD8 localization to ER-mitochondrial MCSs^12,16^, a subset of PDZD8 puncta colocalize with mitochondrial membranes (Fig. 3H and Fig. S3E). Together, our findings are consistent with the conclusion that PDZD8 is enriched at ER-LELs MCSs in neurons.

### PDZD8 promotes autolysosomal turnover

To investigate how PDZD8 promotes autophagy, we quantified autophagic structures labeled by a neuronally expressed mCherry-Atg8a reporter, which marks all autophagic stages^40^, in control and *PDZD8* null mutants. We detect the accumulation of autophagic structures in neuronal cell bodies of *PDZD8* mutants compared to control (Fig. 4A-C). Given PDZD8’s co-localization with LELs, we co-expressed a GFP-Lamp1 reporter^41^ to quantify Atg8a+/Lamp1+ autolysosomes^42^. We detect a significant increase in Atg8a+/Lamp1+ structures in *PDZD8^KO^* (Fig. 4A,C). The accumulation of autolysosomes in *PDZD8* mutants suggests PDZD8 may promote autolysosome turnover. To independently confirm these observations and assess autophagy progression, we used a tandem-tagged mCherry-GFP-Atg8a reporter^43^ (Fig. 4D). Upon transport from autophagosomes to autolysosomes, GFP is rapidly quenched due to the acidic pH, while mCherry persists. Structures positive for both GFP and mCherry (autophagosomes) are not different in *PDZD8^KO^* and controls (Fig. 4E,F white arrowheads). In contrast, structures positive for mCherry alone (autolysosomes) are significantly increased in *PDZD8^KO^*, consistent with stalled autolysosome maturation (Fig. 4E,F magenta arrowheads). These observations predict that ectopic PDZD8 might accelerate autolysosome turnover. We tested this prediction by overexpressing *PDZD8* in neurons together with mCherry-Atg8a and GFP-Lamp1 reporters. Upon *PDZD8* overexpression, we observe a significant decrease in Atg8a+ structures compared to controls (Fig. 4A,C). Consistently, using the tandem-tagged mCherry-GFP-Atg8a reporter, we find that overexpression of P*DZD8* leads to enhanced clearance of autolysosome structures (Fig. 4G,H). These findings indicate that PDZD8 increases autophagic flux and suggest that increased autophagic flux is particularly important during activity-dependent synapse formation.

**Figure 4.**
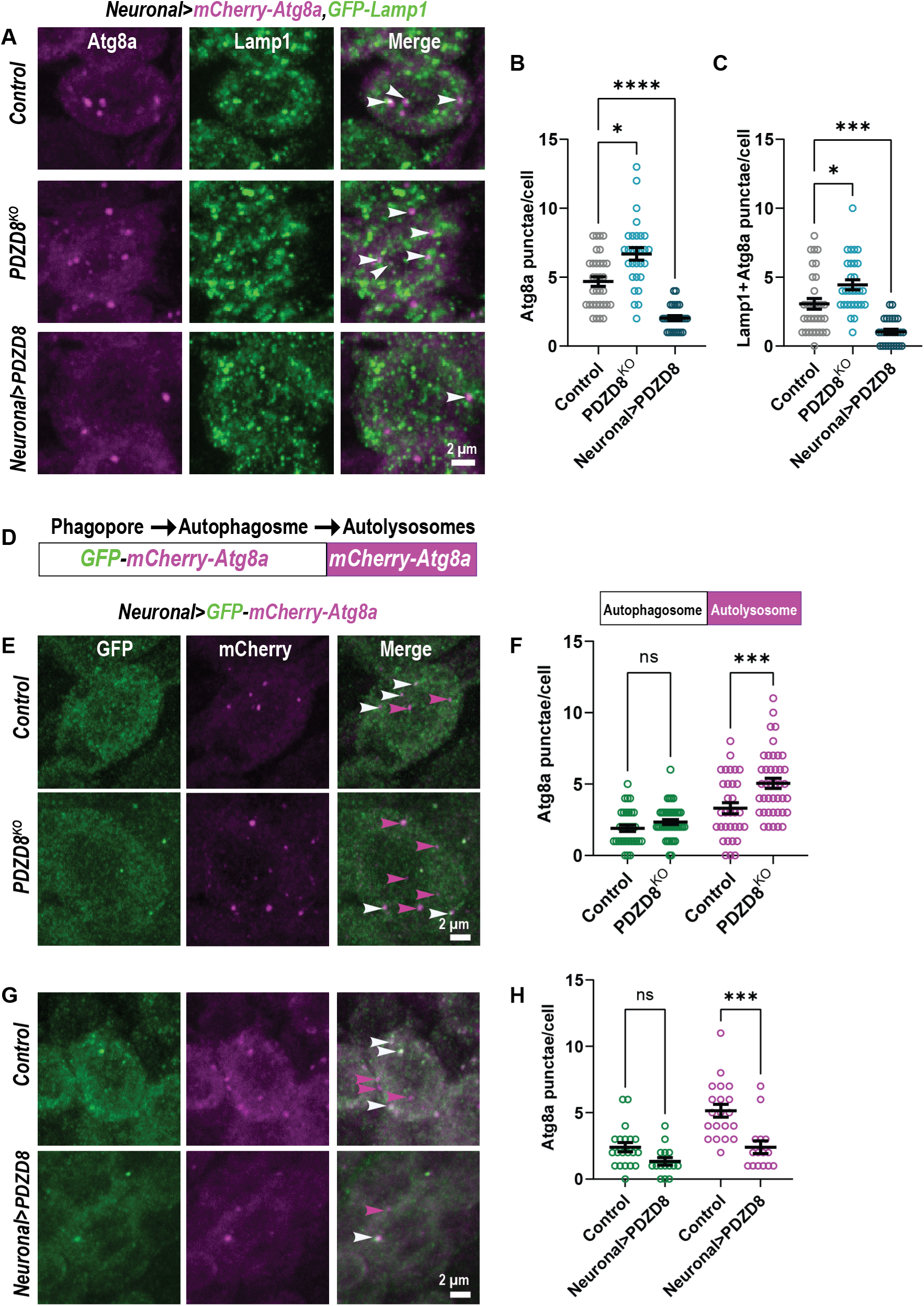
PDZD8 promotes autolysosomal turnover. **(A-C)** PDZD8 regulates autolysosome turnover. Confocal Z-projections of larval neuronal cell bodies co-labeled with antibodies against mCherry (magenta) and GFP (green) **(A)**, quantification of Atg8a puncta **(B)**, and quantification of Lamp1-positive Atg8a punctae **(C)** per cell in the indicated genotypes. White arrowheads indicate Lamp1-positive Atg8a structures. *elav-Gal4* was used to drive expression in neurons. Scale bar=2 μm. **(D-H)** The tandem tagged Atg8a reporter **(D)** distinguishes autophagosomes from autolysosomes and can be used to assess autophagic flux. Confocal Z-projections of larval neuronal cell bodies **(E,F)** co-labeled with antibodies against mCherry (magenta) and GFP (green) and quantification of Atg8a puncate **(G,H)** per cell in the indicated genotypes. White arrowheads point to Atg8a structures positive for both mCherry and GFP (autophagosomes) and magenta arrowheads point to Atg8a structures labeled by mCherry alone (autolysosomes). *elav-Gal4* is used to drive expression in neurons. Scale bar=2 μm. Data is represented as mean ± SEM with significance calculated by ANOVA followed by Tukey’s multiple comparisons test. *p < 0.05,***p < 0.001,****p < 0.0001, n.s. = not significant.

### PDZD8 promotes autolysosome catalytic activity

Given PDZD8’s role as an ER-LEL tether, we hypothesized that PDZD8 promotes lysosome maturation and, thus, autolysosome turnover. The tandem-tag reporter assay indicates that autolysosomes acidify, at least to a degree sufficient to quench GFP, in the absence of PDZD8 (see Fig. 4D-H), so we investigated the hydrolytic activity of lysosomes/autolysosomes. We labeled adult brains with Magic Red, a reporter that fluoresces upon cleavage by lysosomal protease Cathepsin B. Magic Red staining is significantly increased upon *PDZD8* overexpression compared to control (Fig. 5A,B). Because our null allele is marked with DsRed we cannot conduct this assay in loss-of-function alleles, so we investigated levels of autophagy substrate p62. p62 levels inversely correlate with autophagic degradation in flies and mammals and provide a measure of autolysosome function^44–46^. p62 levels are elevated in *PDZD8^KO^* (Fig. 5C,D), consistent with a role for PDZD8 in promoting the degradative activity of autolysosomes.

**Figure 5.**
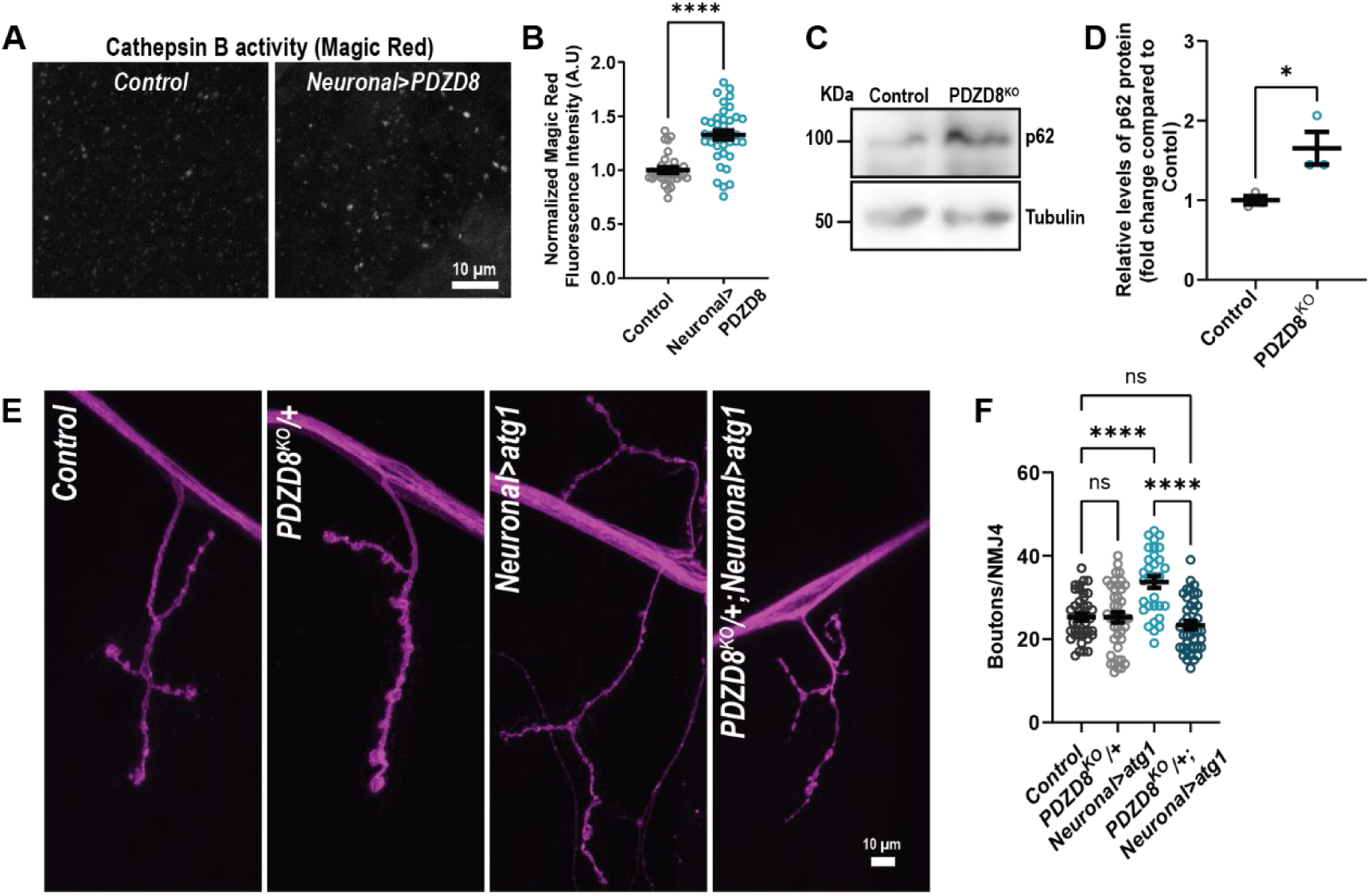
PDZD8 promotes autolysosome catalytic activity. **(A,B)** PDZD8 promotes lysosome proteolytic activity. Representative confocal images of adult brains **(A)** stained with Magic Red cresyl violet-(RR)2 to monitor lysosomal Cathepsin B enzymatic activity and quantification of normalized Magic Red fluorescence intensity per neuron in the indicated genotypes. Same brain regions were used for quantification **(B)**. *elav-Gal4* was used to drive expression in the neurons. Scale bar=10 μm. **(C,D)** Immunoblot of *Drosophila* larval brain extracts labeled with an antibody against p62 **(C)**. Tubulin is used for a loading control. Quantification of fold change in anti-p62 levels relative to WT control **(D)**. **(E,F)** PDZD8 is required for Atg1-induced ectopic synapse formation. Representative confocal images of NMJs labeled with FITC-conjugated anti-HRP (magenta) **(E)** and bouton quantification **(F)** in the indicated genotypes. *elav-Gal4* was used to drive *atg1* expression in neurons. Scale bar=10 μm. Data is represented as mean ± SEM with significance calculated by Mann-Whitney test **(B),** Unpaired t-test **(D)** or ANOVA followed by Tukey’s multiple comparisons test **(F)**. *p < 0.05,****p < 0.0001, n.s. = not significant.

Taken together, our findings lead to the model that PDZD8 promotes autolysosomal maturation to promote autophagic flux and synaptogenesis. Specifically, we propose that PDZD8 is required to accelerate autophagy during periods of high demand such as activity-dependent synaptic growth. If this is the case, decreasing PDZD8 abundance would be expected to suppress the ability of *atg1* overexpression to induce ectopic bouton formation. As previously demonstrated ^24^, we observe significant ectopic bouton formation when we neuronally express *atg1* under the control of the *elav-Gal4* driver. Loss of a single copy of *PDZD8* has no effect in a wild-type background, but completely suppresses Atg1-induced NMJ overgrowth (Fig. 5E,F). These findings further support the model that PDZD8 promotes autolysosomal maturation to increase autophagic flux and promote synaptogenesis.

### PDZD8 localization to ER-LEL MCSs is required to promote autophagy

PDZD8 is a multidomain protein anchored to ER through an N-terminal transmembrane domain. Following the transmembrane domain, is the SMP lipid-transfer domain, a PDZ protein interaction domain, and a C1 lipid-binding domain at the C-terminus^13,14^ (Fig. 6A). While mammalian PDZD8 contains a C-terminal coiled-coil domain, this domain does not appear to be fully conserved at the sequence level in *Drosophila*. Despite this difference in the C terminus, immunoprecipitation mass spectrometry indicates that *Drosophila* PDZD8 still interacts, directly or indirectly, with Rab7 (log2(fold change) increase relative to control = 4.89, FDR adjusted p=0.005), consistent with their co-localization (see Figs 3F, S3C). The C1 domain of mammalian PDZD8 was recently shown to bind a number of phospholipids^13^. To assess the lipid binding ability of the *Drosophila* PDZD8 C1 domain, we incubated strips spotted with phosphatidylinositol phosphate (PIP) lipids with cell extracts of adult heads expressing V5-tagged full-length PDZD8 (PDZD8-V5) and truncated PDZD8 lacking its C terminus (PDZD8^ΔCterm^-V5) in neurons (Fig. S5A). Full-length PDZD8 interacts with phosphatidylserine (PS), phosphatidylinositol (PI) and phosphatidylinositol monophosphates PI3P, PI4P and PI5P (Fig. 6B). With the deletion of the C terminus, PDZD8’s ability to bind lipids is completely lost (Fig. 6B), indicating that *Drosophila* PDZD8 also interacts with lipids through its C terminus. We next explored the significance of lipid binding on the localization of PDZD8 to ER-LEL MCSs *in vivo*. As observed earlier, neuronally expressed PDZD8-V5 localizes ER and ER MCSs similar to endogenously tagged PDZD8 (Fig. 6C and see Fig. 3E-H). However, deletion of the C terminus specifically abolishes PDZD8 localization to LELs without impacting ER localization or localization to ER-mitochondrial MCSs (Fig. 6C and Fig. S5B). Thus, the C-terminus of PDZD8 binds lipids and specifically localizes PDZD8 to ER and LEL MCSs in neurons. This finding allows us to test the functional relevance of ER-LEL MCSs in neurons. To investigate the role of PDZD8 function at ER-LEL MCSs in autophagy, we used the mCherry-Atg8a reporter to quantify overall levels of autophagic structures in neurons overexpressing *PDZD8^ΔCterm^*. Whereas overexpression of full-length *PDZD8* once again accelerates the clearance of autophagic intermediates, overexpression of *PDZD8^ΔCterm^* has no effect (Fig. 6D,E and see Fig. 4A-C). We also investigated the hydrolytic activity of lysosomes/autolysosomes and observed that, unlike full-length PDZD8, overexpression of PDZD8^ΔCterm^ does not lead to increased Magic Red staining (Fig. 6F,G). Together, these findings demonstrate that PDZD8’s lipid-binding C terminus links ER-resident PDZD8 to LELs where it functions to promote lysosome maturation.

**Figure 6.**
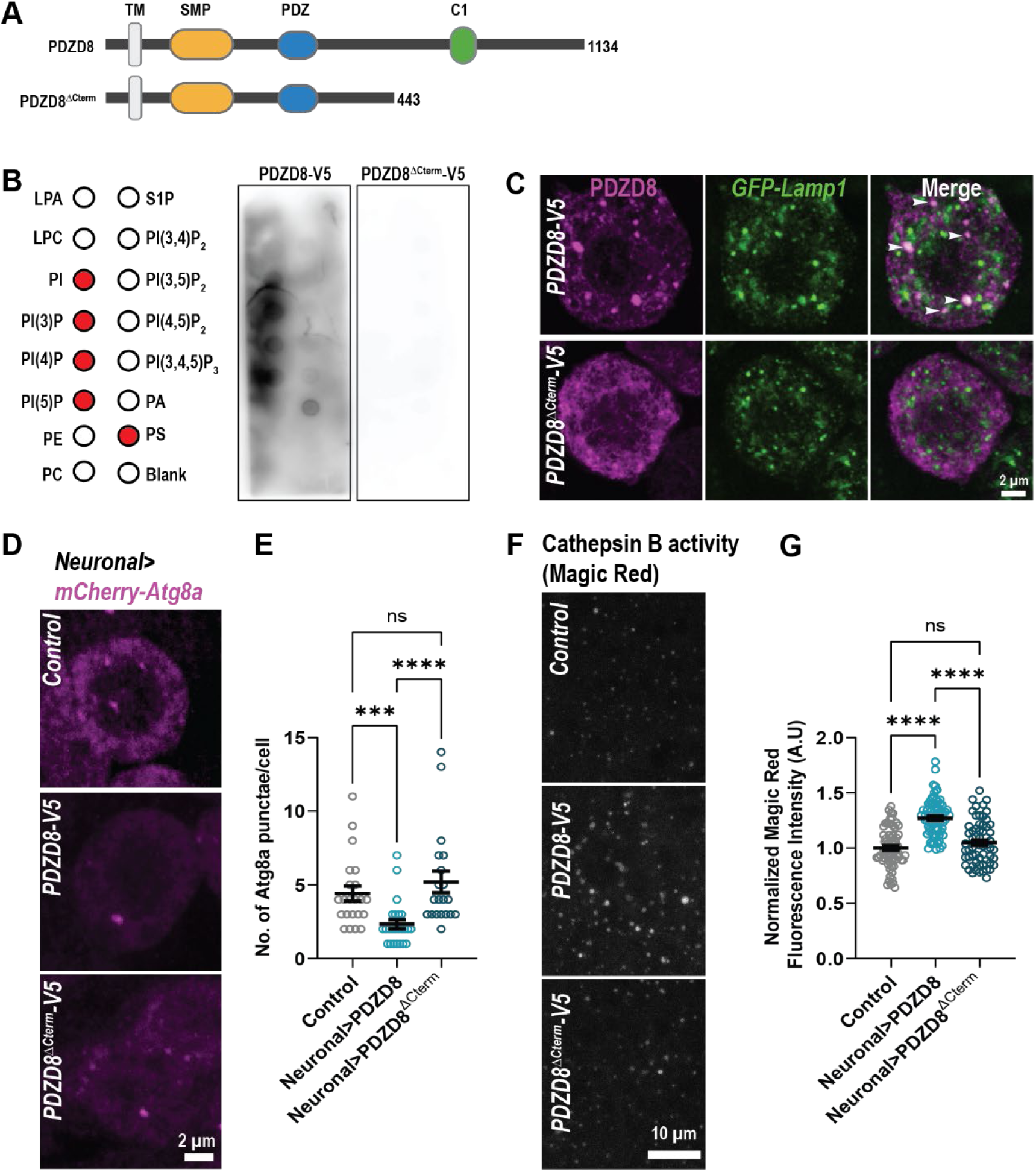
PDZD8 localization to ER-LEL MCSs is required to promote autophagy. **(A)** PDZD8 is a transmembrane (TM) protein with an SMP lipid-transfer domain at the N terminus followed by PDZ protein interaction domain and a C1 lipid-binding domain at the C-terminus. Cartoon depicting the domain organization of full-length PDZD8 and the lipid-binding mutant PDZD8^ΔCterm^ with C terminus deleted. **(B)** PDZD8 interacts with lipids through the C-terminus. Cartoon showing arrangement of lipids on the PIP strip: Lysophosphatidic acid (LPA), Lysophosphocholine (LPC), Phosphatidyinositol (PtdIns), phosphatidylinositol monophosphates PtdIns(3)P, s(4)P, PtdIns(5)P, Phosphatidylethanolamine (PE), Phosphatidylcholine (PC), Sphingosine-1-phosphate (S1P), phosphatidylinositol bisphosphate PtdIns(3,4)P_2_, PtdIns(3,5)P_2_, PtdIns(4,5)P_2_, phosphatidylinositol bisphosphate PtdIns(3,4,5)P_3_, Phosphatidic acid (PA), and Phosphatidylserine (PS). Red circles indicate lipid binding to PDZD8. PIP strips incubated with cellular extracts from *Drosophila* heads expressing full-length *PDZD8-V5* and *PDZD8^ΔCterm^-V5*, respectively, and probed with antibodies against V5. Blots were developed together and imaged simultaneously. *elav-Gal*4 was used to drive *PDZD8* transgene expression in neurons. **(C)** PDZD8 is localized to ER-LELs MCSs through its lipid binding C-terminus. SoRa spinning disk deconvolved single plane of neuronally expressed GFP-Lamp1 in of *elav-Gal4; PDZD8-V5* and *elav-Gal4; PDZD8^ΔCterm^-V5* larval neuronal cell bodies co-labeled with antibodies against GFP (green) and V5 (magenta). Scale bar=2 μm. **(D,E)** PDZD8 localization to ER-LEL MCSs is required to promote autophagy. Confocal Z-projections of larval neuronal cell bodies expressing *mCherry-Atg8* and labeled with antibodies against mCherry (magenta) **(D)** and Atg8a puncta quantification **(E)** in the indicated genotypes. *elav-Gal4* was used to drive expression in neurons. Scale bar=2 μm. **(F,G)** Deletion of PDZD8’s C-terminus impairs lysosomal activity. Representative confocal images of adult brains stained with Magic Red cresyl violet-(RR)2, to monitor lysosomal Cathepsin B enzyme activity **(F)** and quantification of normalized Magic Red fluorescence intensity per cell **(G)** in the indicated genotypes. *elav-Gal4* was used to drive expression in neurons. Scale bar=10 μm. Data is represented as mean ± SEM with significance calculated by Kruskal-Wallis followed by Dunn’s multiple comparisons test **(E,G)**. ***p < 0.001,****p < 0.0001, n.s. = not significant.

### PDZD8 lipid binding and transfer promote synaptic growth

The finding that C-terminus-mediated localization of PDZD8 to ER-LEL MCSs is required for PDZD8’s role in promoting autolysosome clearance allows us to test the functional relevance of this role in promoting synaptic growth. To do this, we compared the ability of full-length *PDZD8* and *PDZD8^ΔCterm^* to induce bouton formation. Whereas neuronal expression of full-length *PDZD8* induces ectopic boutons as expected, overexpression of *PDZD8^ΔCterm^* has no effect on bouton number (Figs. 7A,B). Thus, we conclude that PDZD8 functions at ER-LEL MCSs to promote synaptic growth via autophagy.

**Figure 7.**
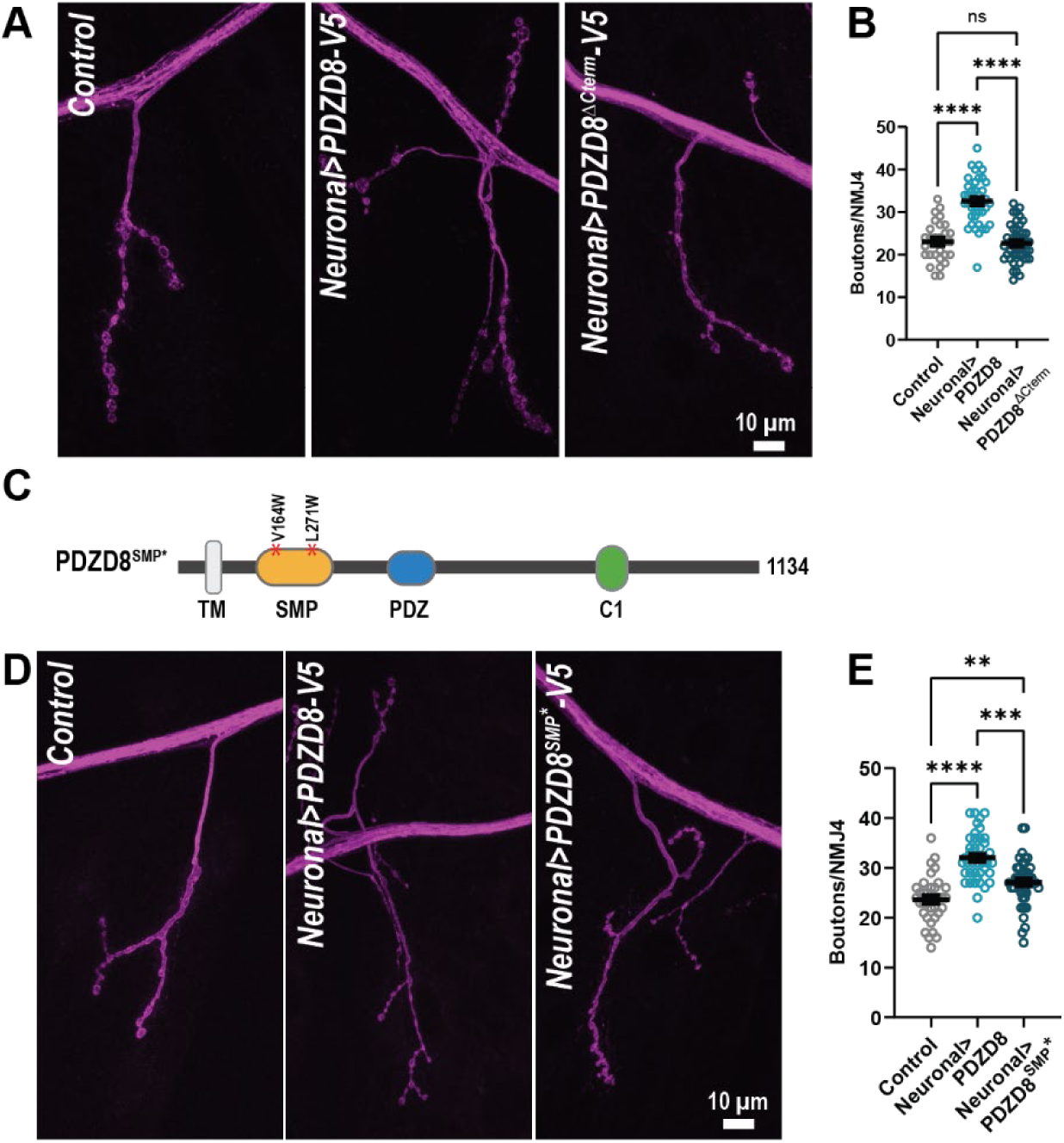
PDZD8 promotes synaptic growth through lipid binding and transfer activity. **(A-B)** ER-LEL MCS localization is critical for PDZD8 induced ectopic synaptic growth. Representative confocal images of NMJs labeled with FITC-conjugated anti-HRP (magenta) **(A)** and bouton quantification **(B)** in the indicated genotypes. *elav-Gal4* was used to drive *PDZD8* transgene expression in neurons. Scale bar=10 μm. **(C-E)** SMP domain mutations significantly impair PDZD8’s ability to promote ectopic synaptic growth. Representative confocal images **(D)** of NMJs labeled with FITC-conjugated anti-HRP (magenta) **(D)** and bouton quantification **(E)** in the indicated genotypes. *elav-Gal4* was used to drive *PDZD8* transgene expression in neurons. Scale bar=10 μm. Data is represented as mean ± SEM with significance calculated by ANOVA followed by Tukey’s multiple comparisons test **(B)** or Kruskal-Wallis followed by Dunn’s multiple comparisons test **(E)**. **p < 0.01,***p < 0.001,****p < 0.0001, n.s. = not significant.

The PDZD8 SMP domain can transport phospholipids *in vitro*^12,13^. To study the functional significance of SMP-dependent lipid transfer activity *in vivo* without disrupting other potential SMP functions ^47^, we replaced two conserved SMP-domain residues containing small hydrophobic side chains with tryptophan residues, which have large side-chains, to generate a space-filling mutant *PDZD8^SMP*^* (V164W, L271W; Fig. 7C). These substitutions were previously shown to strongly impair, but not eliminate, SMP-mediated lipid transfer^12,48^. Neuronally expressed *PDZD8^SMP*^* localizes to ER-LEL and ER-mitochondria MCSs similar to full-length PDZD8 (Fig. S6A-C and see Fig. 3), indicating no requirement for SMP function in localization to MCSs and allowing us to assess the role of SMP-mediated lipid transfer activity in promoting synaptic growth. Overexpression of *PDZD8^SMP*^* in neurons significantly reduces PDZD8’s ability to induce ectopic synaptic bouton formation (Fig. 7D,E). Together, these findings demonstrate *in vivo* roles for PDZD8 lipid binding and transfer activity in promoting synaptogenesis.

## DISCUSSION

The formation and function of synaptic connections requires coordinating a host of cellular activities from transcription to protein trafficking and turnover to energy homeostasis, placing a high demand on cellular communication. This high demand coupled with the polarized structure of neurons and long distances between synapses and the soma present a uniquely challenging environment for intracellular communication. MCS are now recognized as key signaling hubs for coordinating a vast array of cellular activities, and their disruption underlies a number of neurological disorders, including lysosomal storage disorders, hereditary spastic paraplegia, and neurodegeneration, indicating critical roles in the nervous system ^10,49^. Through *in vivo* studies in *Drosophila*, we have found that the MCS tethering protein PDZD8 functions at ER-LEL MCSs to accelerate autophagy during activity-dependent synaptogenesis.

We observe accumulation of autolysosomes in neuronal cell bodies in the absence of PDZD8 and accelerated depletion upon PDZD8 overexpression, suggesting PDZD8 accelerates the clearance of autolysosomes. The accumulation of autophagy cargo protein p62 in PDZD8 mutants further supports a role in promoting autophagic flux. Our findings indicate that PDZD8 functions downstream of autophagosome-lysosome fusion, pointing to a role in maturation. Using a GFP-RFP tandem-tagged Atg8 reporter, which labels autolysosomes with only RFP due to the quenching of GFP in the low-pH environment, we found that autolysosomes acidify at least partially in *PDZD8* mutants. In contrast, we find that overexpression of *PDZD8* enhances lysosomal protease Cathepsin B activity, consistent with a role in promoting the degradative function of autolysosomes. We propose that PDZD8 enhances the degradative capacity of autolysosomes to increase autophagic flux during periods of high demand. Notably, autolysosomes also accumulate in the absence of PDZD8-interactor Rab7 in fed conditions and Rab7 has similarly been proposed to increase autophagic flux by promoting the maturation of lysosomes through the targeting or processing of Cathepsins ^50^. Lipid binding and ER-LEL localization are required for PDZD8’s role in increasing autophagic flux and promoting synaptogenesis, leading us to speculate that PDZD8-dependent lipid transport at MCSs may regulate the lipid composition of lysosomes to promote the accumulation and/or activation of lysosomal proteases^51^. In support, a recent study in *C. elegans* found that PDZD8, plays a role in regulating PI(4,5)P2 accumulation on late endosomes and, thus, cargo degradation ^52^. Future studies to investigate PDZD8’s impact on autolysosome lipid composition and identify the relevant protein targets of PDZD8-mediated autophagy during activity-dependent growth will be of great interest.

Neurons exhibit constitutive and compartmentalized autophagy ^53^. Autophagosomes form in the cell body and at presynaptic terminals. Autolysosomes formed at terminals are retrogradely transported to the soma and mature during transport ^27 25,26,54,55^. We observe an increase in autolysosomes in neuronal soma in the absence of PDZD8. Given PDZD8’s localization in axons and terminals, it will be of interest to determine if PDZD8 plays a role in the maturation of both soma- and synapse-derived autolysosomes.

In addition to these roles at ER-LEL MCSs, PDZD8 has also been shown to function at ER-mitochondria MCSs to regulate dendritic Ca^2+^ dynamics and mitochondrial quality control^11,16^. Interestingly, PDZD8 can form three-way MCSs between ER, LELs and mitochondria ^14^. The hierarchy and the molecular nature of this interaction is unclear. We observe a small fraction of PDZD8 localizing at ER-mitochondrial MCSs in neurons. However, deletion of the C terminus abolishes localization to ER-LEL MCSs without affecting localization to ER-mitochondria MCSs, indicating that PDZD8 recruitment to ER-mitochondria MCSs is independent of its recruitment to ER-LEL MCSs. The observation that disruption of ER-LEL localization completely suppresses PDZD8’s ability to induce synaptic growth further indicates that PDZD8 promotes synaptic growth independently of its role at ER-mitochondria MCSs.

The finding that PDZD8 promotes synaptic growth via autophagy adds to a growing understanding of the importance of neuronal autophagy in synaptic development, function, and plasticity ^56–58^. Autophagy was first shown to promote bouton formation at the *Drosophila* NMJ through negative regulation of Hiw^24^, an E3 ubiquitin ligase that negatively regulates the Wnd/DLK pathway^59^. Here, we expand on this work to define a role for autophagy in the formation of individual motor synapses as well. PDZD8’s role in basal bouton or synapse formation at the NMJ is only revealed when autophagy is limiting, indicating it is not required for, but rather enhances or accelerates, autophagy. We hypothesize that robust autophagy is particularly important during periods of rapid synaptogenesis. Notably, *Drosophila PDZD8* transcription is highly spatio-temporally restricted to periods of embryonic/larval and pupal synaptogenesis^60^. Recent studies in the *Drosophila* visual system underscore the importance of temporally regulated autophagy during circuit formation ^28,29^. We also found that PDZD8 is required for activity-dependent synaptic growth when the requirement for autophagy is likely high as increased synaptic activity has been shown to induce autophagy at the *Drosophila* NMJ^34 61^. In this context, PDZD8-dependent acceleration of autophagy could serve as a mechanism for translating neural activity into synaptic growth.

Vamp-Associated Proteins (VAPs) are broad mediators of ER MCSs^62^. At the *Drosophila* NMJ, VAP-33A may also link autophagy to synaptic growth. Loss of *VAP-33A* results in fewer synaptic boutons, whereas overexpression induces ectopic formation of small boutons, similar to both *PDZD8* and *atg1* ^24,63^. Recent studies reveal roles for VAPA/B in promoting autophagosome biogenesis at contact sites between ER and isolation membranes (the first step in autophagosome formation) and VAP33 in autolysosome acidification at ER-Golgi MCSs^64,65^. Autophagy also promotes synaptic growth in *C. elegans*, where notably it has distinct roles in different neurons^30^. In one neuronal subtype, Stavoe et al., found that autophagy regulated axon outgrowth. Studies in neuronal culture suggest that PDZD8 may also regulate axon outgrowth in some neurons^13 12,66^. The regulatory diversity of autophagy in synapse formation is also revealed by the observation that in some contexts autophagy promotes synaptogenesis ^18,24,30,31^, while in others it attenuates synapse formation^28,29,32^. These findings point to context-specific roles for autophagy in sculpting synaptic connections to promote neural circuit formation and function. While autophagy’s critical role in neurodegenerative disorders is well established, growing links between regulators of autophagy, including PDZD8, and neurodevelopmental disorders underscore the importance of expanding our understanding of its role in nervous system development and plasticity^67^.

## Supplemental Information

**Figure S1.**
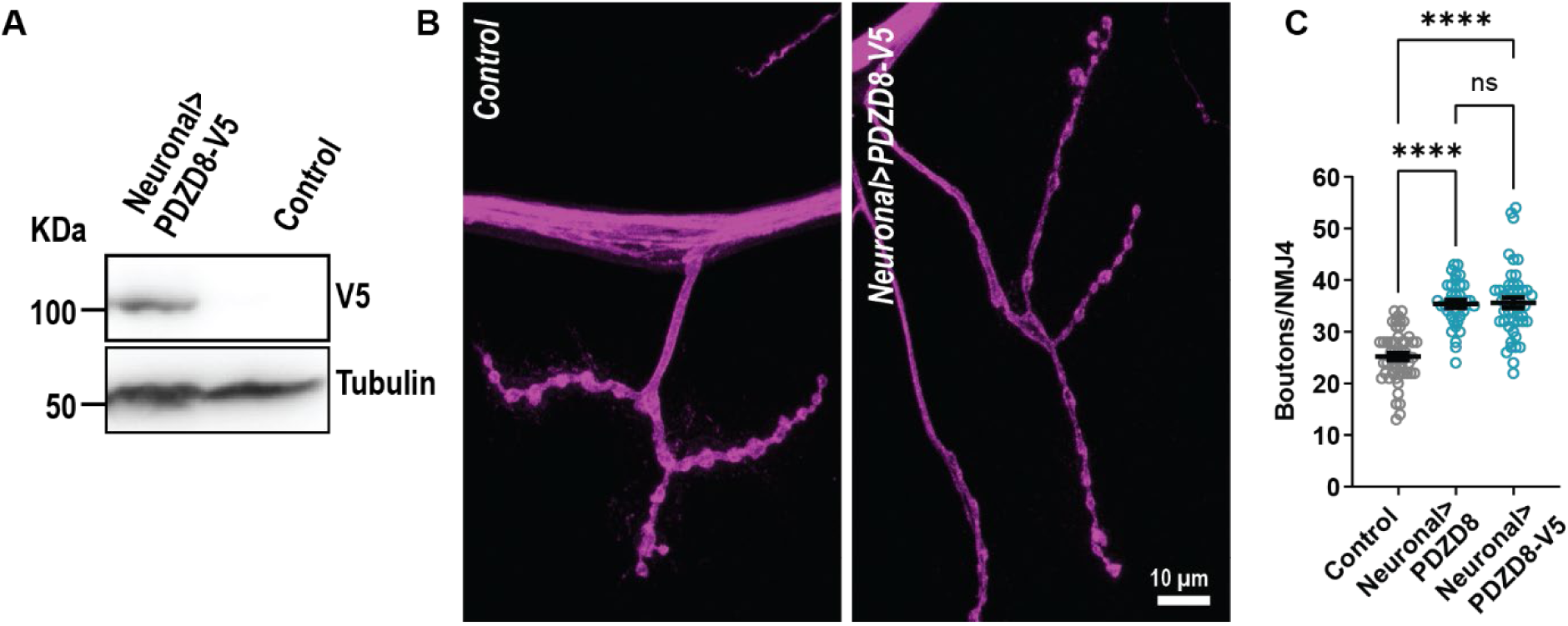
C-terminal tagging does not impair PDZD8 function. **(A-C)** Neuronal over expression of *PDZD8-V5* induces ectopic synaptic growth similar to untagged *PDZD8*. Immunoblot of *Drosophila* head extracts **(A)** probed with anti-V5 antibody demonstrate the stable expression of PDZD8-V5 protein. Tubulin is used as loading control. Representative confocal images of NMJs labeled with FITC-conjugated anti-HRP (magenta) **(B)** and bouton quantification **(C)** in the indicated genotypes. *elav-Gal4* was used to drive *PDZD8* expression in neurons. The data for Control and Neuronal>PDZD8 are the same as Fig. 1F. Scale bar=10 μm. Data is represented as mean ± SEM with significance calculated by Kruskal-Wallis followed by Dunn’s multiple comparisons test. ****p < 0.0001, n.s. = not significant.

**Figure S2.**
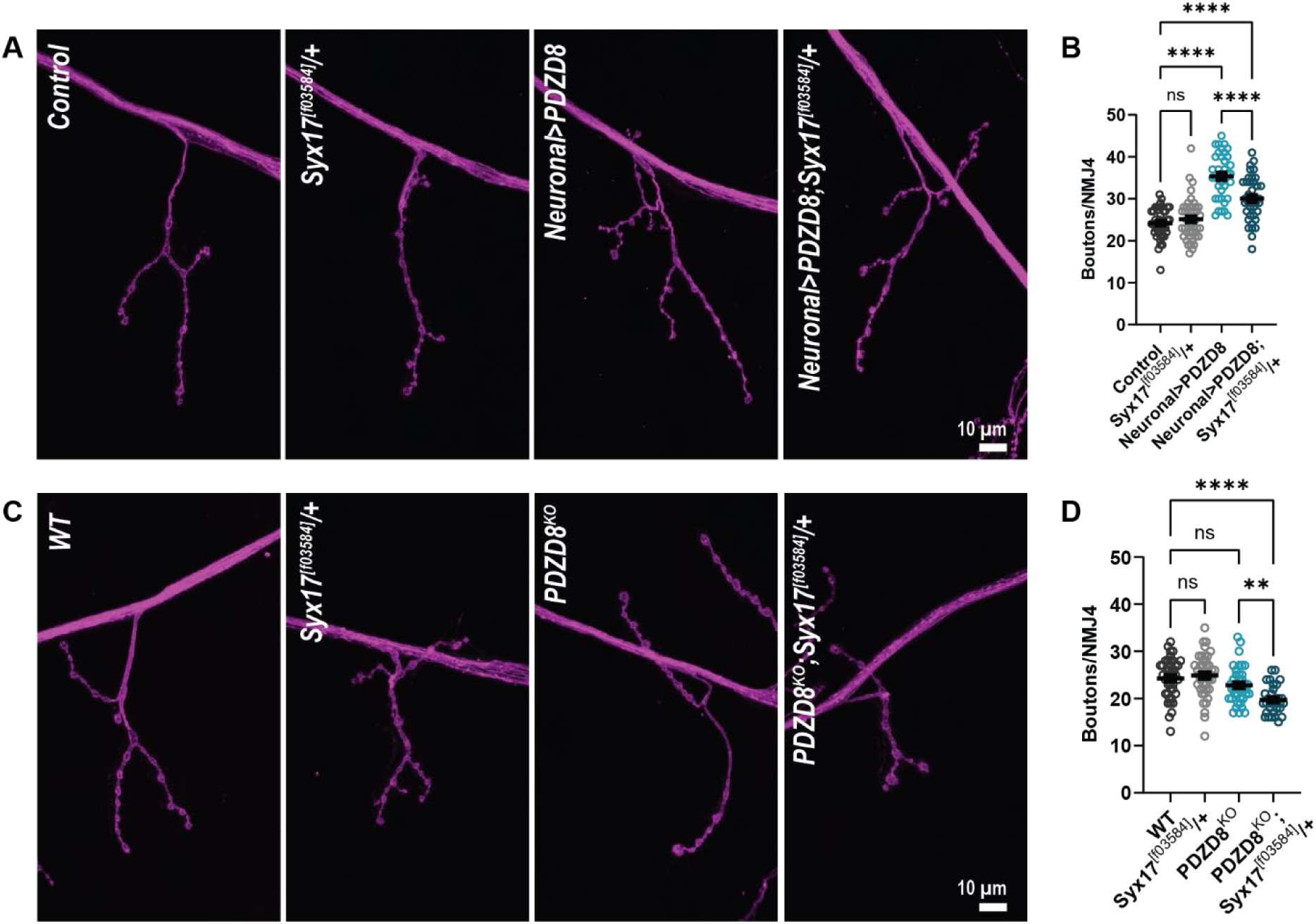
PDZD8 and syntaxin 17 genetically interact. **(A-D)** Downregulation of autophagy by reducing the levels of Syx17 suppresses PDZD8-induced ectopic synaptic growth and dominantly enhances loss of *PDZD8 l*eading to basal synaptic undergrowth. Representative confocal images labeled with FITC-conjugated anti-HRP (magenta) **(A,C)** and bouton quantification **(B,D)** in the indicated genotypes. *elav-Gal4* was used to drive *PDZD8* expression in the neurons. Scale bar=10 μm. Data is represented as mean ± SEM with significance calculated by or ANOVA followed by Tukey’s multiple comparisons test **(B,D)**. **p < 0.01,****p < 0.0001, n.s. = not significant.

**Figure S3.**
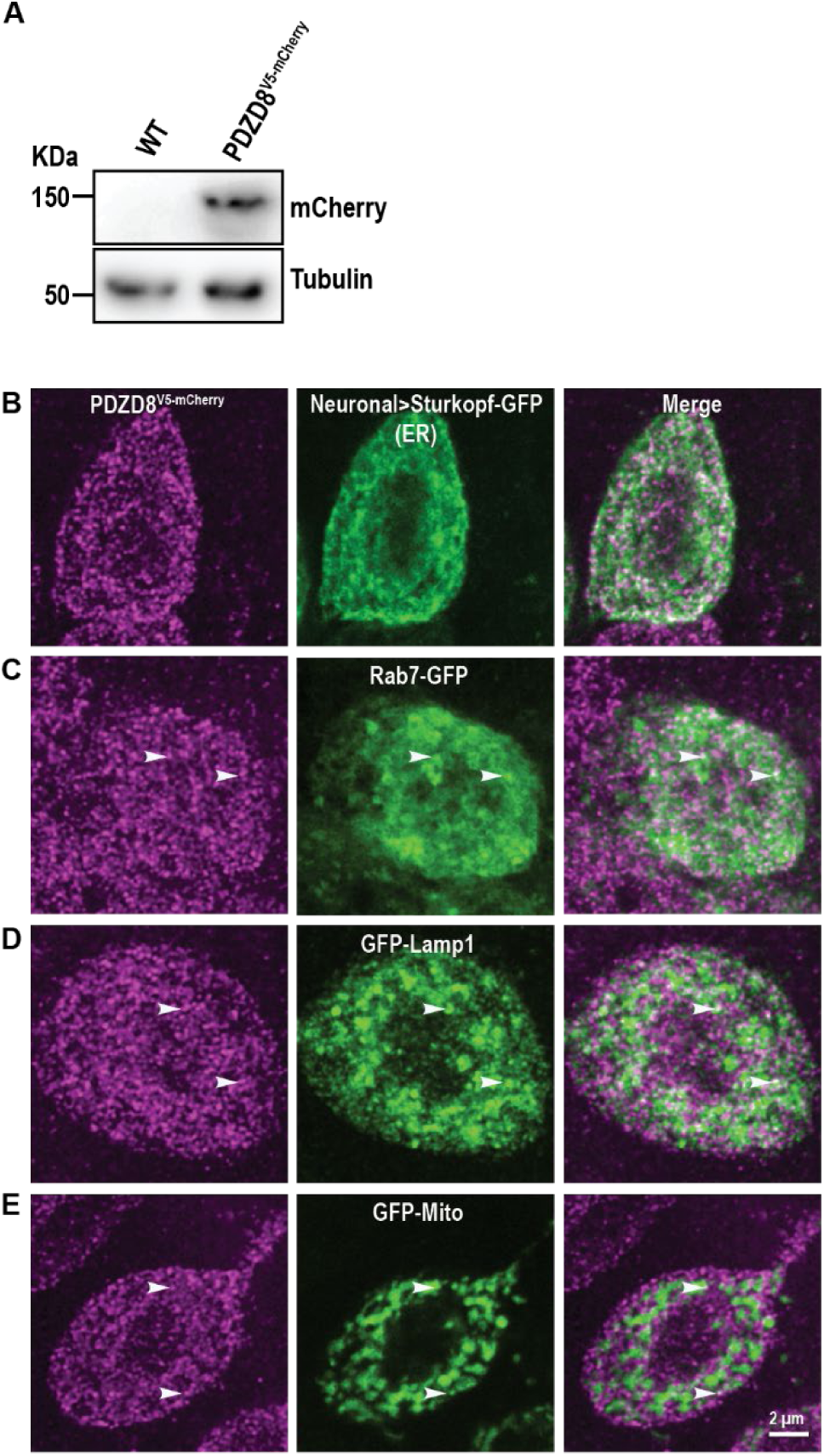
Endogenous PDZD8 localizes to ER-LEL membrane contact sites in neurons. **(A)** Immunoblot of *Drosophila* head extracts probed with anti-mCherry antibody demonstrates stable expression of endogenously tagged PDZD8 protein of the expected size. Tubulin is used as a loading control. **(B-E)** SoRa spinning disk deconvolved single Z-stack of a *PDZD8^V^*^5^*^-mCherry^* larval neuronal cell body co-labeled with antibodies against mCherry (magenta) and in green Sturkopf-GFP to label ER **(B)**, Rab7-GFP to label late endosomes **(C)**, GFP-Lamp1 to label lysosomes**(D)**, and GFP-Mito to label mitochondria **(E).** White arrowheads indicate colocalization **(B-E)** *OK6-Gal4* was used to drive transgene expression in neurons. Scale bar=2 μm.

**Figure S4.**
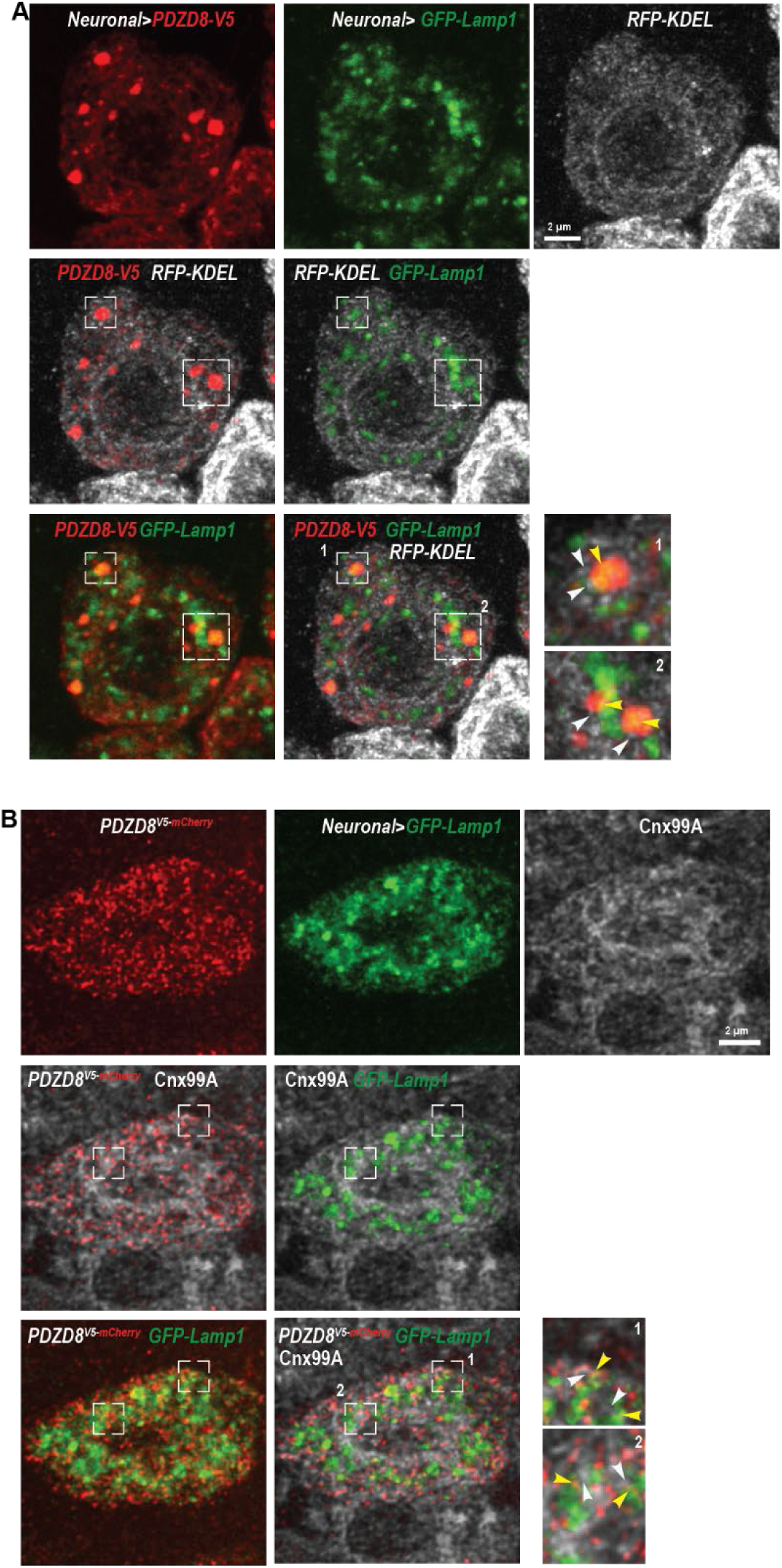
PDZD8 localizes to ER-Lysosomes MCSs. **(A,B)** PDZD8 localizes to ER-Lysosomes MCS. SoRa spinning disk deconvolved single Z-stack of larval neuronal cell body of PDZD8-V5 **(A)** and PDZD8^V5-mCherry^ **(B)** co-labeled with antibodies against V5 or mCherry (red) respectively, GFP (green) labels the lysosomes with GFP-Lamp1 and RFP-KDEL or Cnx99A (white) labels the endoplasmic reticulum. White square boxes indicate the region of colocalization of PDZD8 with ER-Lysosomes MCSs. Boxes 1 and 2 are zoomed-in to show the co-localization of PDZD8 and Lamp1 (yellow arrow head) with KDEL (white arrow head). Elav-Gal4 is used to drive expression of PDZD8-V5 and endomembrane markers in the neurons whereas OK6-Gal4 is used to drive GFP-Lamp1 in PDZD8^V5-mCherry^. Scale bar-2 μm.

**Figure S5.**
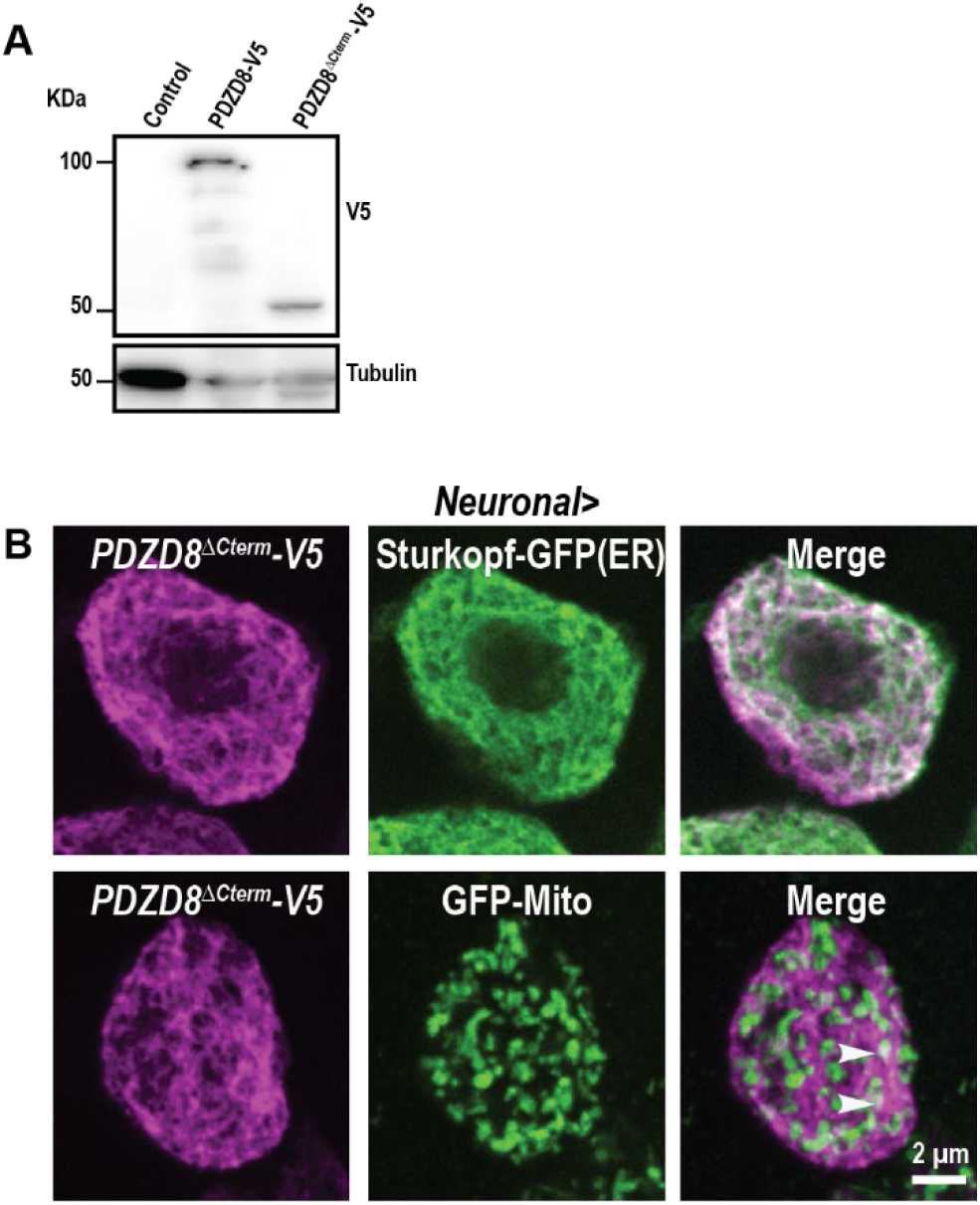
C-terminal-mediated lipid binding is dispensable for PDZD8 localization to ER-mitochondria MCSs. **(A,B)** PDZD8^ΔCterm^-V5 is stably expressed and produces a protein of predicted size. Immunoblot of *Drosophila* head extracts probed with anti-V5 antibody **(A)** Tubulin is used as a loading control. SoRa spinning disk deconvolved single Z-stack of larval neuronal cell body of PDZD8^ΔCterm^-V5 co-labeled with antibodies against V5 (magenta) and GFP (green) to label ER with Sturkopf-GFP and mitochondria with GFP-Mito **(B)**. White arrowheads indicate sites of colocalization of PDZD8 and mitochondria. *elav-Gal4* was used to drive expression in neurons. Scale bar=2 μm.

**Figure S6.**
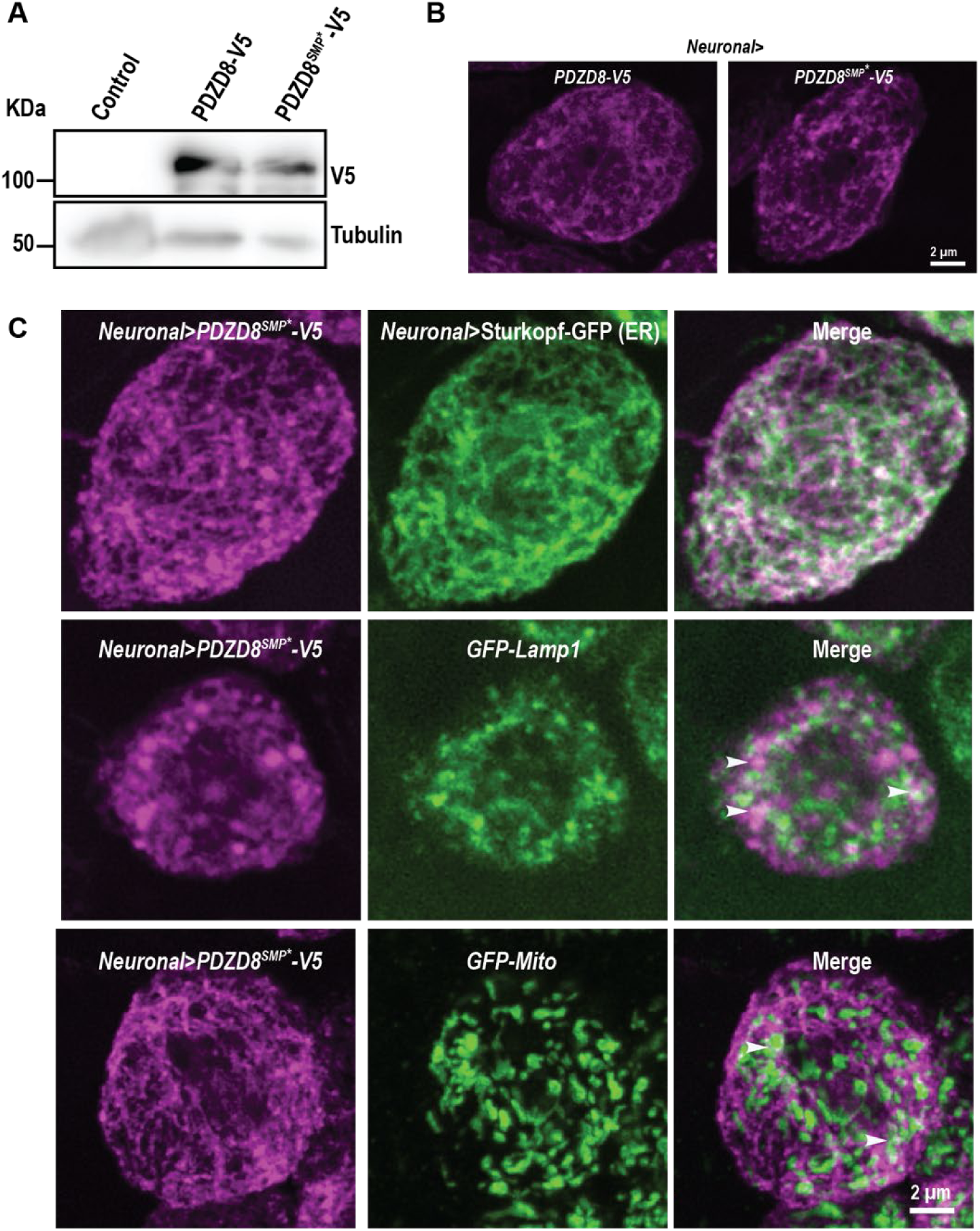
Expression and localization of PDZD8^SMP*^-V5. **(A-C)** Introduced mutations in the PDZD8 SMP domain do not affect stable expression or sub-cellular localization. Immunoblot of *Drosophila* head extracts probed with anti-V5 antibodies **(A)** demonstrates that WT-PDZD8 and PDZD8^SMP*^are expressed similarly. Tubulin is used as a loading control. PDZD8 ^SMP*^localizes similarly to full-lengthPDZD8. SoRa spinning disk deconvolved single Z-stack of larval neuronal cell bodies of PDZD8-V5 and PDZD8 ^SMP*^-V5 labeled with antibodies against V5 (magenta) **(B)** and co-labeled with antibodies against GFP (green) **(C)** to label ER with Sturkopf-GFP, lysosomes with GFP-Lamp1, and mitochondria with GFP-Mito. White arrowheads indicate sites of colocalization of PDZD8 with lysosomes or mitochondria. *elav-Gal4* was used to drive expression in neurons. Scale bar=2 μm.

## STAR METHODS

### KEY RESOURCES TABLE

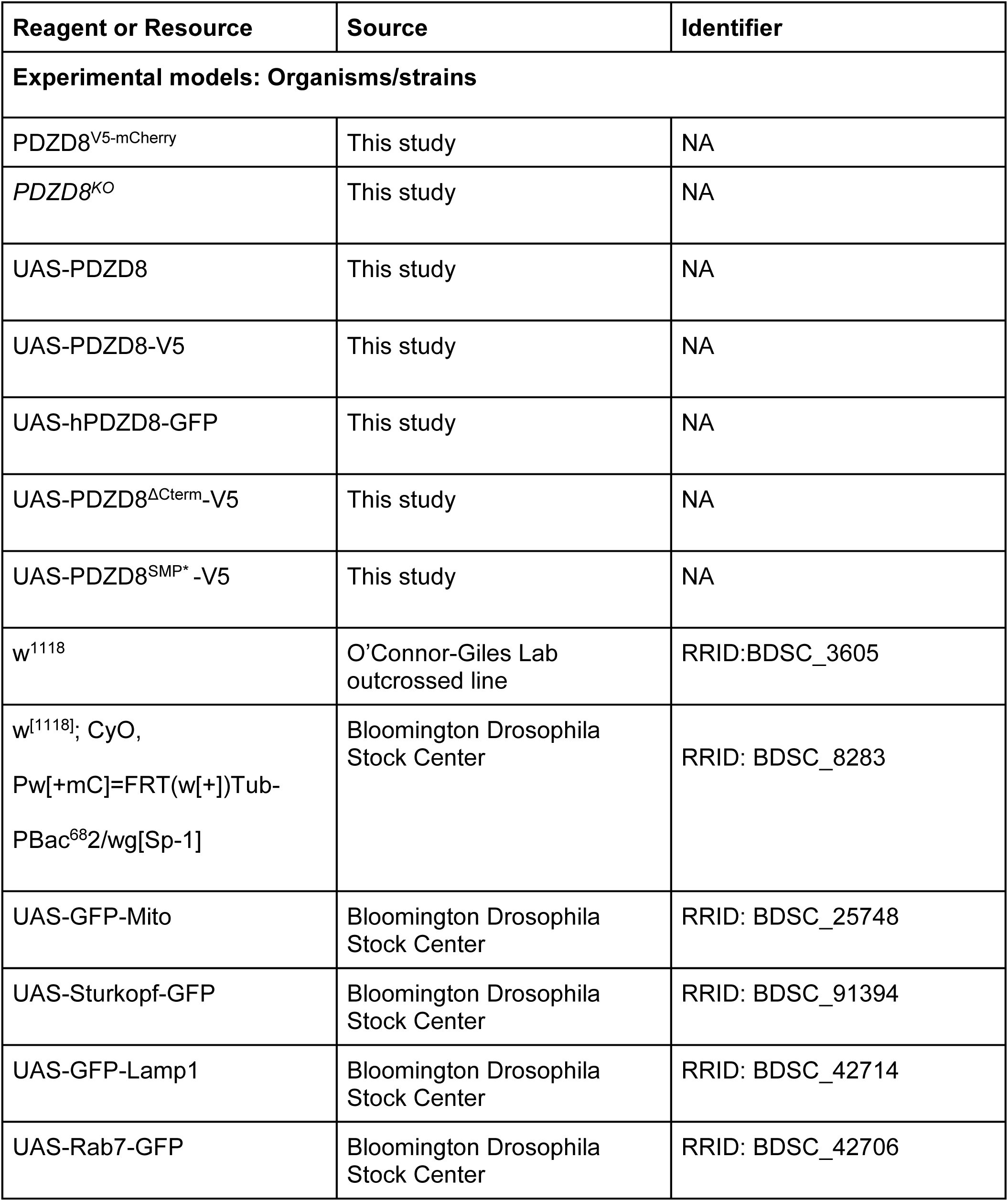
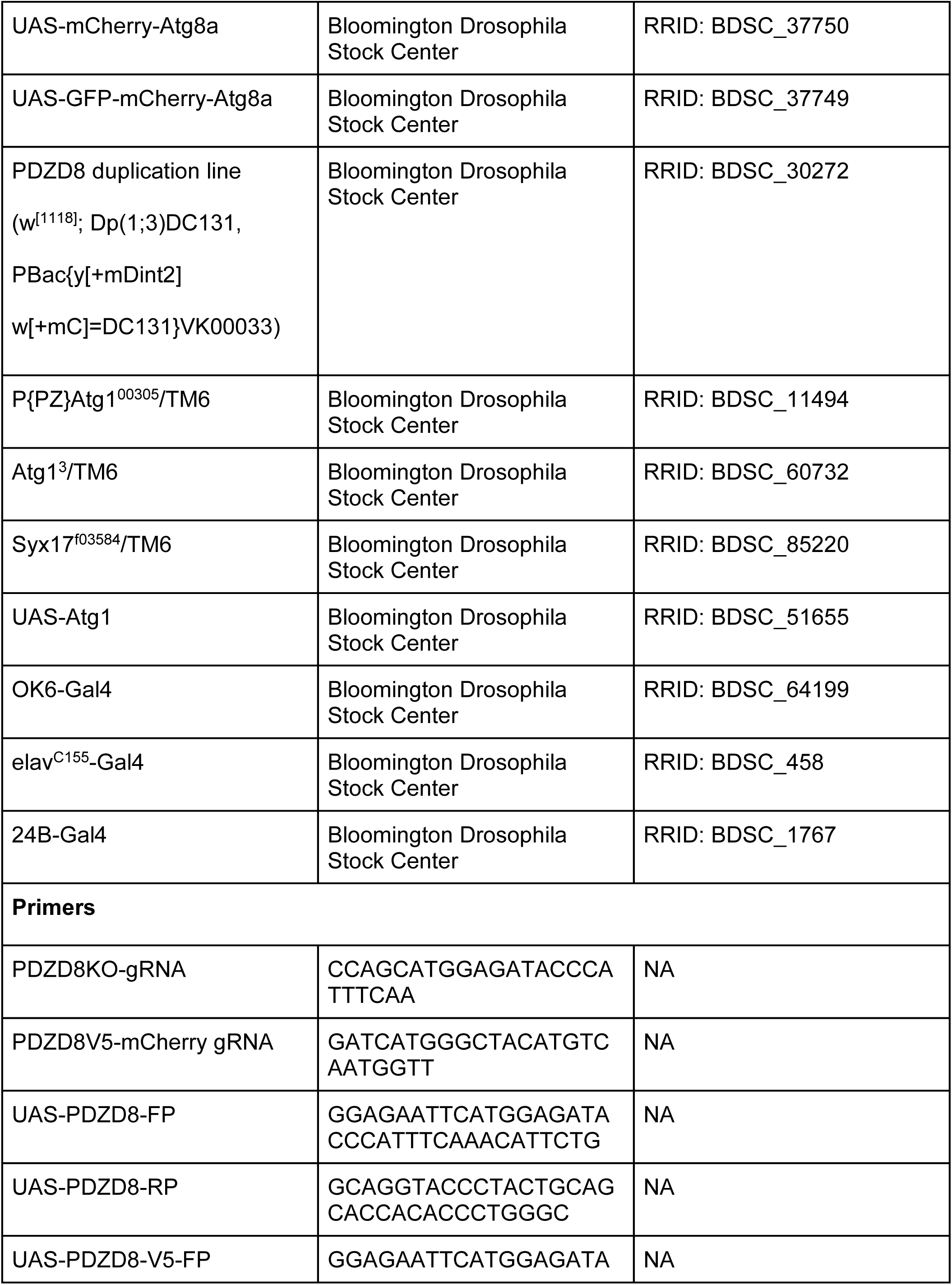
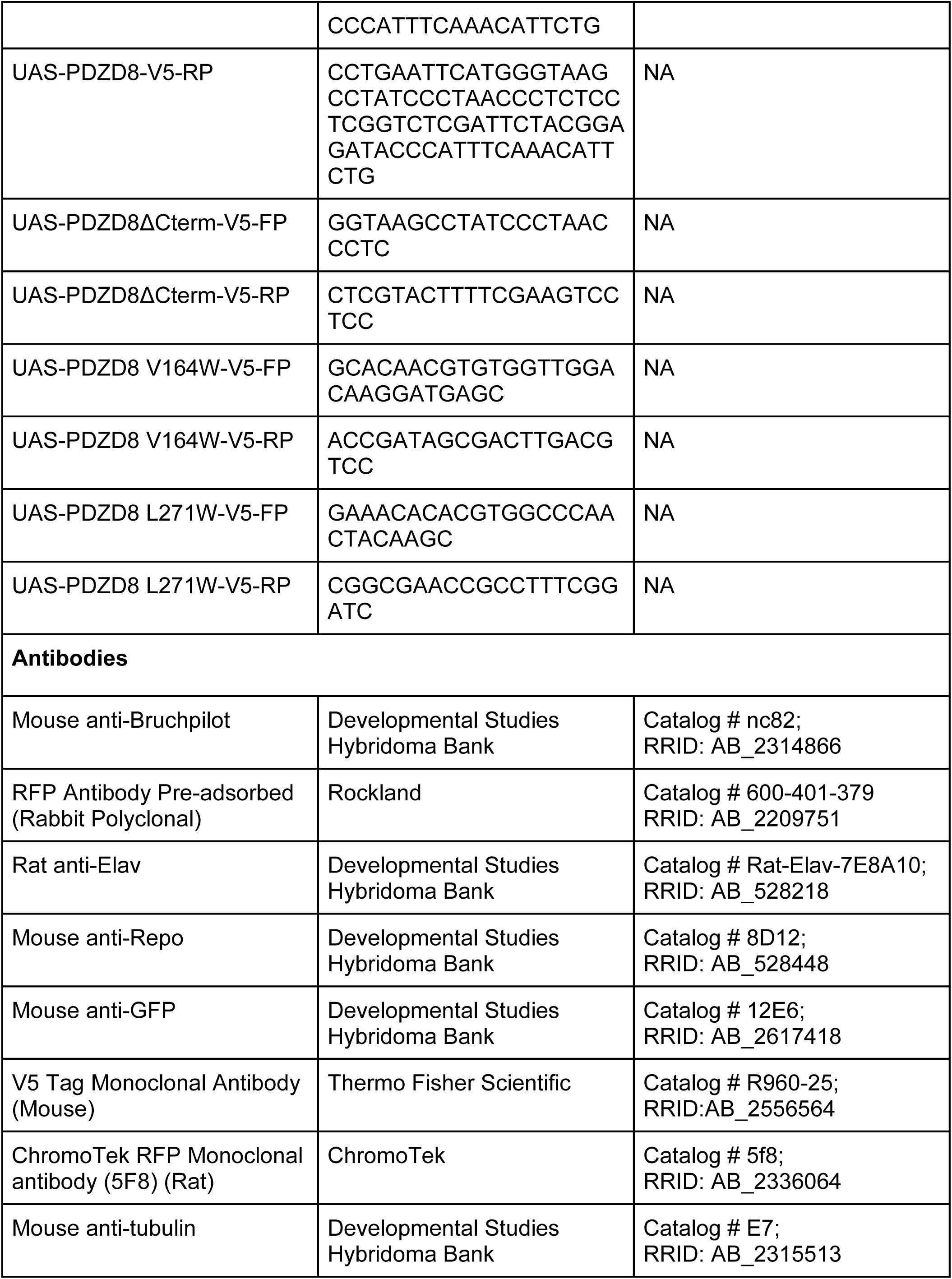
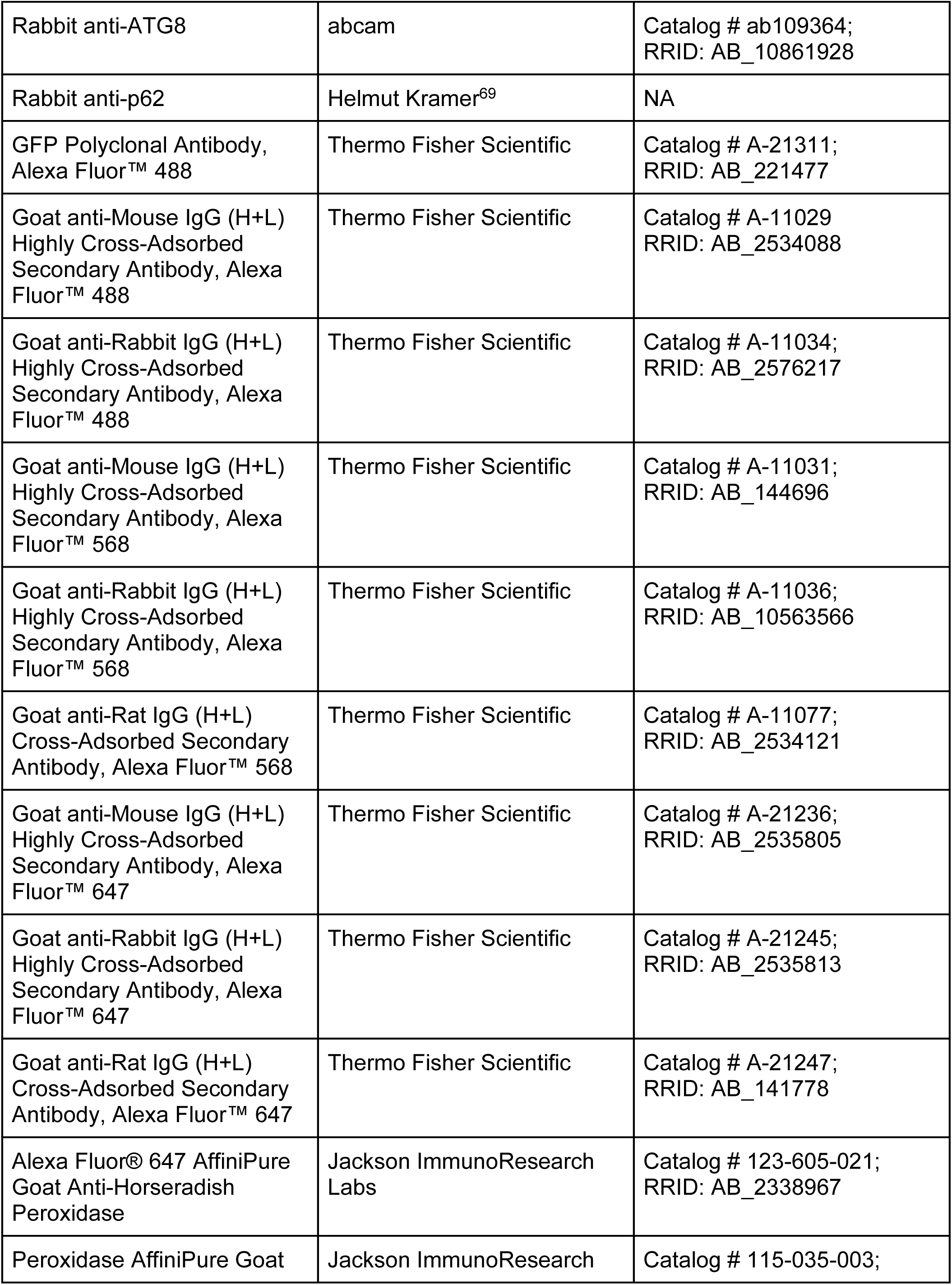
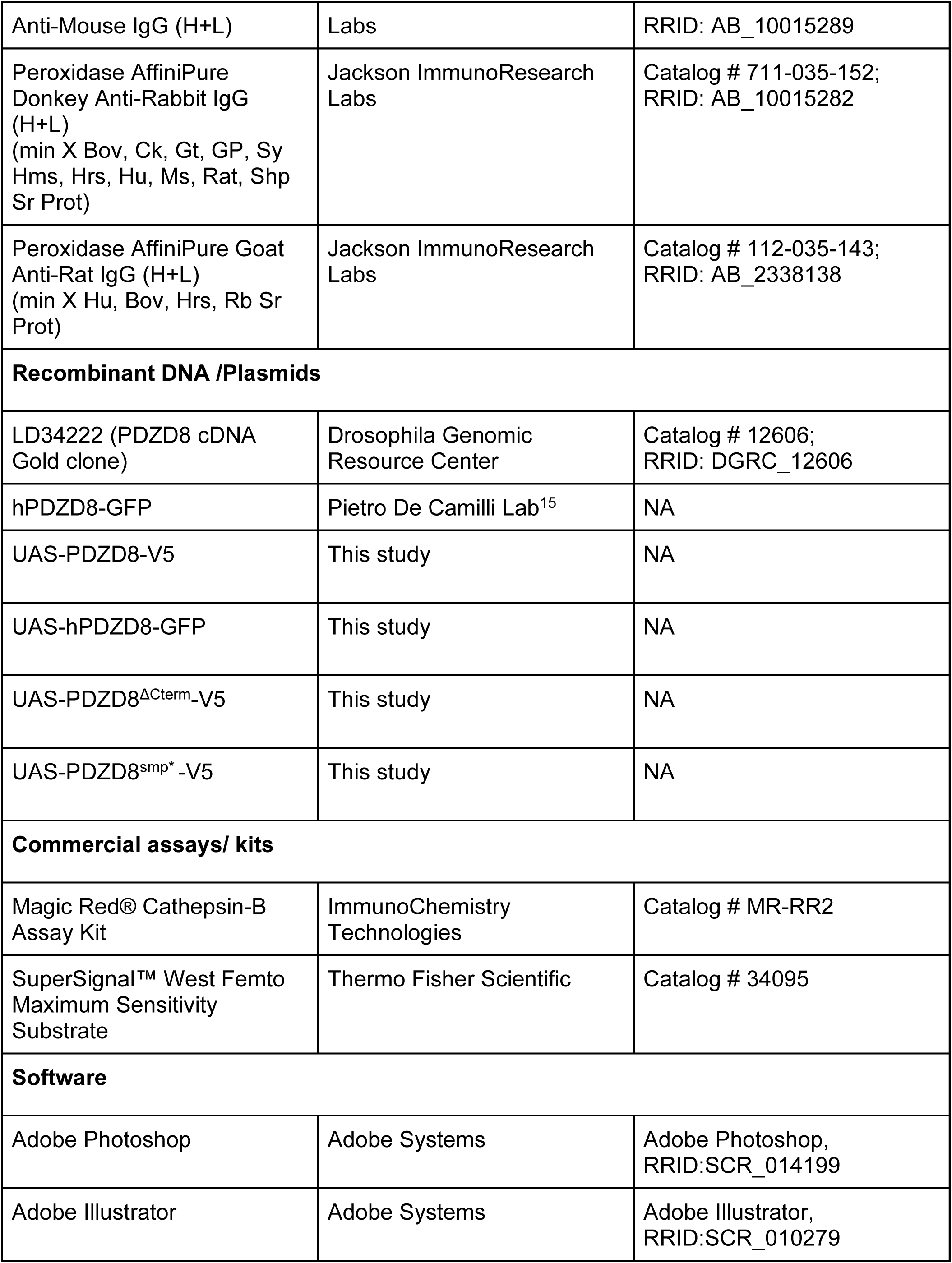
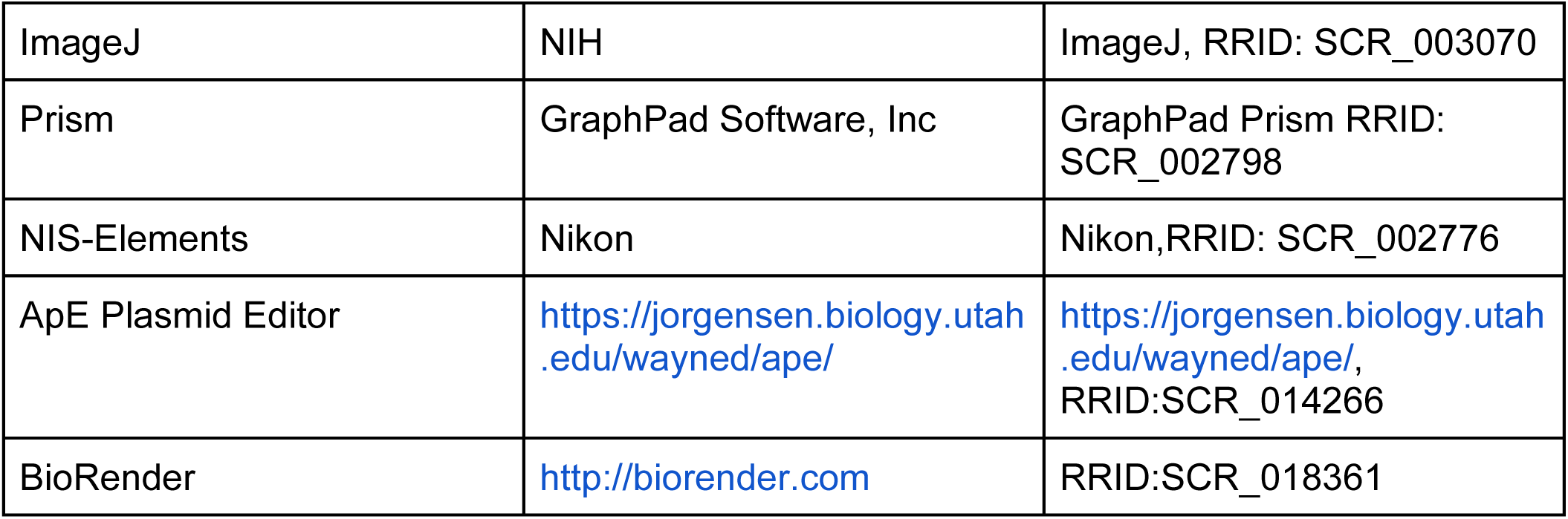

### RESOURCE AVAILABILITY

#### Lead contact

Requests for any resources or reagents should be addressed to the lead contact, Kate M. O’Connor-Giles.

#### Materials availability

All plasmids, transgenic flies, and custom reagents created for this study are available upon request to the lead contact.

### EXPERIMENTAL MODEL AND SUBJECT DETAILS

#### Drosophila stocks

All control genotypes, mutant combinations, and crosses were maintained on cornmeal medium (Fly food R, Lab Express, Ann Arbor, MI) at 25°C and 60% relative humidity with a 12/12 light/dark cycle in specialized incubators (Darwin Chambers, St. Louis, MO). *w*^1118^ (BDSC#5905) was used as a WT control, unless otherwise noted. The following lines were acquired from the Bloomington Drosophila Stock Center: *UAS-GFP-mCherry-Atg8a* (BDSC#37749), *UAS-mCherry-Atg8a* (BDSC#37750), *UAS-GFP-Lamp1* (BDSC#42714), *UAS-GFP-mito* (BDSC#25748), *UAS-Rab7-Rab7* (BDSC#42706), *UAS-Sturkopf-GFP* (BDSC#91394), *PDZD8*-duplication line-*w*[1118]*; Dp(1;3)DC131, PBac{y[+mDint2] w[+mC]=DC131}VK00033* (BDSC#30272), *P{PZ} atg1^00305^*/TM6 (BDSC#11494), *atg1^3^*/TM6 (BDSC#60732), *UAS-Atg1* (BDSC#51655), *Syx17*^*f*03584^/TM6 (BDSC#85220), *OK6-Gal4* (BDSC#64199), *elav^C155^–Gal4* (BDSC#458) and *24B-Gal4* (BDSC#1767).

The following genetic lines were generated in this study: *PDZD8^V^*^5^*^-mCherry^*, *PDZD8^KO^*, *UAS-PDZD8*, *UAS-PDZD8-V5*, *UAS-hPDZD8-GFP*, *UAS-PDZD8^ΔCterm^* and *UAS-PDZD8^SMP*^*.

### METHOD DETAILS

#### Generation of endogenously tagged and null alleles of PDZD8

A CRISPR-based homology directed repair strategy was used to generate the following in-frame endogenously tagged and null alleles:. gRNA target sites were selected using the CRISPR Optimal Target Finder program (http://targetfinder.flycrispr.neuro.brown.edu; ^70^). gRNA and donor plasmids were generated as described in Bruckner et al., 2017^71^ and at flycrispr.com. *Vasa-Cas9* embryos were injected with a mixture ofone gRNA plasmids (100ng/μL, each) and a double-stranded DNA donor plasmid (500ng/μL) by BestGene, Inccrossed to *w*^1118^ flies after eclosion, and progeny screened for DsRed expression in the eye. All alleles were edited using the scarless CRISPR-piggyBac approach (flycrispr.com; (Bruckner et al., 2017)). Briefly, a peptide or protein tag flanked by flexible linkers and followed by a visible marker flanked by piggyBac inverted terminal repeat sequences was inserted immediately downstream of the sole *PDZD8* translational start site. This generates a null allele due to interruption of the open reading frame and the presence of stop codons in the visible marker cassette. By crossing edited lines to piggyBac (BL8285) transposase, the visible marker cassette can be removed to generate in-frame tags. For the endogenous C-terminal V5-mCherry tag, the same approach was used to insert the tag immediately before the sole stop codon. All engineered lines were confirmed by Sanger sequencing of the locus.

#### Generation of UAS lines

UAS transgenes were generated by cloning full-length or truncated *Drosophila*(Drosophila Genomics Resource Center #1261) or human (provided by Andrés Guillén-Samander and Pietro De Camilli) *PDZD8* cDNAs into pUAST-C5. All transgenes were integrated into the attP2 landing site on the third chromosome (BDSC#8622) by BestGene, Inc. Site-directed mutagenesis was used to introduce the V164W and L271W mutations in UAS-PDZD8^SMP*(V164W,L271W)^.

#### Immunostaining and confocal imaging

Third instar larvae were dissected in Ca^2+^-free saline and fixed for 6 min in Bouin’s fixative, except for *PDZD8^V^*^5^*^-mCherry^*, which was fixed in ice cold methanol for 5 min on ice. Dissected larvae were washed and permeabilized in PBS with 0.1% Triton-X, blocked in PBS containing 0.1% Triton-X, 5% NGS and 1% BSA for 30 min at room temperature (RT) or overnight at 4℃, followed by overnight incubation with primary antibodies and 4-hour incubation with secondary antibodies at room temperature, then mounted in Vectashield (Vector Laboratories). For superresolution imaging, samples were mounted in ProLong™ Glass Antifade Mountant (ThermoFisher Scientific #P36980) and cured overnight. The following primary antibodies were used at the indicated concentrations: Mouse anti-Bruchpilot at 1:100 (Developmental Studies Hybridoma Bank #NC82,), Rat anti-Elav at 1:100 (Developmental Studies Hybridoma Bank #7E8A10), Mouse anti-GFP at 1:100 (Developmental Studies Hybridoma Bank #12E6), Mouse anti-Repo at 1:25 (Developmental Studies Hybridoma Bank # 8D12), Rabbit anti-RFP at 1:500 (Rockland # 600-401-379), Rat anti-RFP at 1:200 (ChromoTek #5F8), and Mouse anti-V5 at 1:500 (ThermoFisher Scientific # R960-25). The following Alexa conjugated primary and secondary antibodies were used: Rabbit anti-GFP Alexa Fluor 488 at 1:500 (ThermoFisher Scientific, #A-21311), Anti-HRP conjugated to Cy3 or Alexa Fluor 647 at 1:500 (Jackson ImmunoResearch Laboratories, Inc), and Alexa Fluor 488/568/647 at 1:500 (ThermoFisher Scientific). Images were acquired on a Nikon A1R HD confocal microscope with a Plan-Apo 60X 1.49 NA oil-immersion objective at a pixel resolution of 130 nm. Super-resolution images were acquired on a Nikon CSU-W1 SoRa (Spinning Disk Super Resolution by Optical Pixel Reassignment) with a Photometrics Prime BSI sCMOS camera and a 60x 1.49 NA oil-immersion objective. Images were acquired using Nikon NIS and deconvolved using Richardson-Lucy deconvolution with 15-20 iterations. Image analysis was performed using Image J ^72–74^ and NIS-Elements software. Primary and Secondary antibody details are listed in the key resource table.

#### Magic Red Staining

To monitor Cathepsin B activity, dissected adult brains were incubated for 5 min in 1x PBS containing 1x Magic Red cresyl violet-(RR)2 (Immunochemistry Technology, Bloomington, MN) per manufacturer’s instructions and imaged immediately on a Nikon A1R HD confocal microscope with a plan-Apo 40X 1.30 NA oil-immersion objective. Similar brain regions were selected for imaging in all samples.

#### Western blotting

Protein samples were prepared from wandering third instar larval nervous systems or 1-day-old fly heads in 2X Laemmli buffer followed by boiling at 95°C for 5 min. Primary antibodies were incubated overnight at 4°C. The following antibodies were used: rabbit anti-ATG8 (1:1000, abcam# ab109364), rabbit anti-p62 (1:5000, generously provided by Helmut Kramer^69^), anti-rabbit RFP (1:1000, Rockland 600 401 379), mouse anti-tubulin (1:4000, Developmental Studies Hybridoma Bank #E7c), and mouse anti-V5 (1:5000,ThermoFisher Scientific). All HRP-conjugated secondary antibodies (Jackson Immunochemicals) were used at 1:10000 dilutions and incubated for 2 hours at room temperature. Blots were developed using SuperSignal™ West Femto Maximum Sensitivity Substrate (Thermo Fisher Scientific) on Azure c600 biosystems. Primary and Secondary antibody details are listed in the key resource table.

#### Lipid-binding analysis

The protocol used was modified from Munnik.et.al for *Drosophila* heads ^75^. Approximately 200 heads were prepared in 150µl of fat blot buffer (FBB,50 mM Tris/Cl-pH 7.5, 150mM NaCl, and freshly added 30ul PIC). Protein extract was centrifuged at 8000 rpm for 15 min at 4°C twice to remove all debris. Total volume was brought to 1 ml with FBB containing 5% BSA. PIP lipid strips (P-6001, Echelon Biosciences) were blocked with 5% BSA in FBB for 1 hour followed by incubation with protein extracts overnight @4°C on a nutator. The blots were washed 5 times with 0.1% Tween-20 in FBB for 10 minutes followed by primary and secondary antibody incubation in FBB containing 0.1% Tween-20 FBB and 5% BSA for 3 hours at RT. Blots were washed 5 times with FBB containing 0.1% Tween-20 for 10 minutes and developed using an SuperSignal™ West Femto Maximum Sensitivity Substrate (Thermo Fisher Scientific; Catalog # 34095).

#### Replication of results

All experiments were carried out in at least three biological replicates, and all crosses were set up at least twice to obtain reproducible results from replicate to replicate.

### QUANTIFICATION AND STATISTICS

All quantifications were conducted masked to genotype and/or treatment.

#### Quantification of synaptic bouton number

Boutons formed by motor neuron 4-Ib on muscle 4 of segments A2, A3 and A4 were labeled with HRP and quantified manually from confocal images. To analyze the number of AZs per NMJ 4, ROIs were drawn using HRP staining. Nikon Elements Software was used to process images using Gaussian and rolling ball filters and the Brightspots module was used for segmentation to accurately identify and count Brp spots (AZs).

#### Quantification of Atg8 and Lamp1 punctae

For Atg8 and or Lamp1 puncta quantification, cell bodies of the larger larval midline motor neurons were analyzed. NIS-Element General Analysis (GA3) was used to perform the analysis. ROIs were manually drawn to mark individual cells in the VNC. Briefly, images were processed using Gaussian and rolling ball filters and the Brightspots module was used for segmentation to accurately identify and count the total number of puncta per cell.

#### Quantification of Cathepsin B activity

Quantification was performed using Image J software^72–74^. 3D images were z-projected to obtain 2D images in Image J. Multiple ROI per image were drawn to measure mean Magic Red intensity across genotypes. The data was normalized to average mean of *Control* Magic Red intensity.

#### Quantification of western blots

Quantification was performed using Image J software^72–74^. The mean background intensities were subtracted from the images. ROIs were drawn around the bands of interest and the integrated intensity of each ROI was extracted from the image. The loading control bands were analyzed in the same manner. Further numerical analyses were performed in Microsoft Excel.

#### Statistical analysis

All graphing and statistical analyses were completed using GraphPad Prism 10. Normally distributed datasets were analyzed using either an unpaired two-sided t-test (two groups) or a one-way ANOVA with Tukey’s multiple comparisons (more than two groups) and non-normally distributed datasets were analyzed using either a two-sided Mann–Whitney test (two groups) or a Kruskal–Wallis test with Dunn’s multiple comparisons (more than two groups). Error bars report ± SEM. P values and statistical tests used are reported in each figure legend and sample sizes are reported in Table 1.

**Table 1:** Full genotypes and sample size for each experiment.

| Figure | Genotypes | N |
| --- | --- | --- |
| **Figure 1** |  |  |
| Figure 1A-B | *W<sup>1118</sup>*<br>*PDZD8<sup>KO</sup>* | 41 NMJs from 8 animals (25°C) and 37 NMJs from 8 animal (31°C)<br>50 NMJs from 8 animals (25°C) and 38 NMJs from 8 animal (31°C) |
| Figure 1C-D | *W<sup>1118</sup>*<br>*PDZD8<sup>KO</sup>*<br>*PDZD8<sup>KO</sup>::; Dp(1;3)DC131, PBac{y[+mDint2] w[+mC]=DC131}VK00033/+ (Rescue)* | 34 NMJs from 8 animals<br>34 NMJs from 8 animals<br>38 NMJs from 8 animals |
| Figure 1E-F | *elav<sup>C155</sup>-Gal4/Y (Control)*<br>*elav<sup>C155</sup>-Gal4/Y;;UAS-PDZD8/+* | 52 NMJs from 10 animals<br>36 NMJs from 8 animals |
| Figure 1G-H | *+/Y;;24B-Gal4 (Control)* | 26 NMJs from 6 animals |
|  | *+Y;;24B-Gal4/UAS-PDZD8* | 34 NMJs from 8 animals |
| Figure 1I | *elav<sup>C155</sup>-Gal4/Y;;UAS-PDZD8-V5/UAS-hPDZD8-Gfp* |  |
| Figure 1J-K | *elav<sup>C155</sup>-Gal4/Y (Control)* | 36 NMJs from 8 animals |
|  | *elav<sup>C155</sup>-Gal4/Y;;UAS-hPDZD8-Gfp/+* | 38 NMJs from 8 animals |
| **Figure 2** |  |  |
| Figure 2A-B | *elav<sup>C155</sup>-Gal4/Y (Control)* | 44 NMJs from 8 animals |
|  | *Atg1<sup>PZ</sup>/+* | 41 NMJs from 8 animals |
|  | *elav<sup>C155</sup>-Gal4/Y;;UAS-PDZD8/+* | 41 NMJs from 8 animals |
|  | *elav<sup>C155</sup>-Gal4/Y;;UAS-PDZD8/Atg1<sup>PZ</sup>* | 40 NMJs from 8 animals |
| Figure 2C-E | *W<sup>1118</sup>* | 40 NMJs from 8 animals |
|  | *Atg1<sup>PZ</sup>/+* | 43 NMJs from 8 animals |
|  | *PDZD8<sup>KO</sup>* | 47 NMJs from 8 animals |
|  | *PDZD8<sup>KO</sup>/Y;;Atg1<sup>PZ</sup>/+* | 43 NMJs from |
|  |  | 8 animals |
| Figure 2F-H | *W<sup>1118</sup>*<br>*Atg1<sup>3/PZ</sup>* | 21 NMJs from<br>4 animals<br>20 NMJs from<br>4 animals |
| **Figure 3** |  |  |
| Figure 3A-D | *PDZD8<sup>V5-mCherry</sup>* |  |
| Figure 3E | *elav<sup>C155</sup>-Gal4/Y;;UAS-PDZD-V5/ UAS-Sturkopf-GFP* |  |
| Figure 3F | *elav<sup>C155</sup>-Gal4/Y;;UAS-PDZD-V5/ UAS-GFP-Lamp1* |  |
| Figure 3G | *elav<sup>C155</sup>-Gal4/Y;;UAS-PDZD-V5/ UAS-Rab7-GFP* |  |
| Figure 3H | *elav<sup>C155</sup>-Gal4/Y;;UAS-PDZD-V5/ UAS-GFP-Mito* |  |
| **Figure 4** |  |  |
| Figure 4A-C | *elav<sup>C155</sup>-Gal4/Y;UAS-mCherry-Atg8a/+;*<br>*UAS-GFP-Lamp1/+ (Control)*<br>*PDZD8<sup>KO</sup>/Y;UAS-mCherry-Atg8a/+;*<br>*UAS-GFP-Lamp1/+*<br>*elav<sup>C155</sup>-Gal4/Y;UAS-mCherry-Atg8a/+;*<br>*UAS-GFP-Lamp1/UAS-PDZD8-V5* | 32 cell bodies<br>from 4 animals<br>29 cell bodies<br>from 4 animals<br>27 cell bodies<br>from 4 animals |
| Figure 4E-F | *elav<sup>C155</sup>-Gal4/Y;UAS-Gfp,mCherry-Atg8a/+* | 22 cell bodies from 4 animals |
|  | *PDZD8<sup>KO</sup>,elav<sup>C155</sup>-Gal4/Y;UAS-GFP, mCherry-Atg8a/+;* | 24 cell bodies from 4 animals |
| Figure 4G-H | *elav<sup>C155</sup>-Gal4/Y;UAS-GFP,mCherry-Atg8a/+* | 22 cell bodies from 4 animals |
|  | *elav<sup>C155</sup>-Gal4/Y;UAS-GFP,mCherry-Atg8a/+; UAS-PDZD8-V5/ +* | 24 cell bodies from 4 animals |
| **Figure 5** |  |  |
| Figure 5A-B | *elav<sup>C155</sup>-Gal4/Y* | 30 cell bodies from 4 animals |
|  | *elav<sup>C155</sup>-Gal4/Y;;UAS-PDZD8-V5/ +* | 37 cell bodies from 4 animals |
| Figure 5E-F | *elav<sup>C155</sup>-Gal4/+ (Control)* | 40 NMJs from 8 animals |
|  | *PDZD8<sup>KO</sup>/+* | 41 NMJs from 8 animals |
|  | *elav<sup>C155</sup>-Gal4/+;;UAS-ATG1/+* | 31 NMJs from 8 animals |
|  | *PDZD8<sup>KO</sup>,elav<sup>C155</sup>-Gal4/+;;UAS-ATG1* | 42 NMJs from 8 animals |
| **Figure 6** |  |  |
| Figure 6B | *elav<sup>C155</sup>-Gal4/Y;;UAS-PDZD8-V5/+*<br>*elav<sup>C155</sup>-Gal4/Y;;UAS-PDZD8<sup>ΔC1-CC</sup>-V5/+* |  |
| Figure 6C | *elav<sup>C155</sup>-Gal4/Y;;UAS-PDZD8-V5/UAS-Gfp-Lamp1*<br>*elav<sup>C155</sup>-Gal4/Y;;UAS-PDZD8<sup>ΔC1-CC</sup>-V5/ UAS-Gfp-Lamp1* |  |
| Figure 6D-E | *elav<sup>C155</sup>-Gal4/Y;UAS-mCherry-Atg8a/+*<br>*elav<sup>C155</sup>-Gal4/Y;UAS-mCherry-Atg8a/+; UAS-PDZD8-V5/ +*<br>*elav<sup>C155</sup>-Gal4/Y;UAS-mCherry-Atg8a/+; UAS-PDZD8<sup>ΔC1-CC</sup>-V5/ +* | 22 cell bodies<br>from 4 animals<br>24 cell bodies<br>from 4 animals<br>20 cell bodies<br>from 4 animals |
| Figure 6F-G | *elav<sup>C155</sup>-Gal4/Y*<br>*elav<sup>C155</sup>-Gal4/Y;;UAS-PDZD8-V5/ +*<br>*elav<sup>C155</sup>-Gal4/Y;;UAS-PDZD8<sup>ΔC1-CC</sup>-V5/ +* | 70 cell bodies<br>from 6 animals<br>77 cell bodies<br>from 6 animals<br>60 cell bodies<br>from 5 animals |
| **Figure 7** |  |  |
| Figure 7A-B | *elav<sup>C155</sup>-Gal4/Y*<br>*elav<sup>C155</sup>-Gal4/Y;;UAS-PDZD8-V5/ +* | 29 NMJs from<br>8 animals<br>41 NMJs from<br>8 animals |
|  | *elav<sup>C155</sup>-Gal4/Y;;UAS-PDZD8<sup>ΔC1-CC</sup>-V5/ +* | 43 NMJs from 8 animals |
| Figure 7D-E | *elav<sup>C155</sup>-Gal4/Y* | 36 NMJs from 8 animals |
|  | *elav<sup>C155</sup>-Gal4/Y;;UAS-PDZD8-V5/ +* | 43 NMJs from 8 animals |
|  | *elav<sup>C155</sup>-Gal4/Y;;UAS-PDZD8<sup>SMP\*</sup>-V5/ +* | 45 NMJs from 8 animals |
| **Supplementary Figure S1** |  |  |
| Supplementary Figure S1B-C | *elav<sup>C155</sup>-Gal4/Y (Control)* | 52 NMJs from 10 animals |
|  | *elav<sup>C155</sup>-Gal4/Y;;UAS-PDZD8-V5/+* | 47 NMJs from 10 animals |
| **Supplementary Figure S2** |  |  |
| Supplementary Figure S2A-B | *W<sup>1118</sup>* | 42 NMJs from 8 animals |
|  | *Syx17<sup>f03584</sup>/+* | 46 NMJs from 8 animals |
|  | *elav<sup>C155</sup>-Gal4/Y;;UAS-PDZD8/+* | 38 NMJs from 8 animals |
|  | *elav<sup>C155</sup>-Gal4/Y;;UAS-PDZD8/Syx17<sup>f03584</sup>* | 39 NMJs from 8 animals |
| Supplementary Figure S2C-D | *W<sup>1118</sup>* | 40 NMJs from 8 animals |
|  | *Syx17<sup>f03584</sup>/+*<br>*PDZD8<sup>KO</sup>*<br>*PDZD8<sup>KO</sup>/Y;;UAS-PDZD8/Syx17<sup>f03584</sup>* | 43 NMJs from 8 animals<br>47 NMJs from 8 animals<br>43 NMJs from 8 animals |
| **Supplementary Figure S3** |  |  |
| Supplementary Figure S3A | *PDZD8<sup>V5-mCherry</sup>* |  |
| Supplementary Figure S3B | *PDZD8<sup>V5-mCherry</sup>/Y;OK6-Gal4/+;UAS-Sturkopf-Gfp/+* |  |
| Supplementary Figure S3C | *PDZD8<sup>V5-mCherry</sup>/Y;OK6-Gal4/UAS-Gfp-Lamp1* |  |
| Supplementary Figure S3D | *PDZD8<sup>V5-mCherry</sup>/Y;OK6-Gal4/+;UAS-Rab7-Gfp/+* |  |
| Supplementary Figure S3E | *PDZD8<sup>V5-mCherry</sup>/Y;OK6-Gal4/+;UAS-Gfp-Mito/+* |  |
| **Supplementary Figure S4** |  |  |
| Supplementary Figure S4A | *elav<sup>C155</sup>-Gal4/Y;UAS-GFP-Lamp1/+; UAS-PDZD8-*<br>*V5/UAS-RFP-KDEL* |  |
| Supplementary Figure S4B | *PDZD8<sup>V5-mCherry</sup>/Y;OK6-Gal4/UAS-GFP-Lamp1* |  |
| **Supplementary Figure S5** |  |  |
| Supplementary Figure S6B | *elav<sup>C155</sup>-Gal4/Y;;UAS-PDZD8<sup>ΔC1-CC</sup>-V5/ UAS-*<br>*Sturkopf-GFP* |  |
|  | *elav<sup>C155</sup>-Gal4/Y;;UAS-PDZD8<sup>ΔC1-CC</sup>-V5/ UAS-GFP-Mito* |  |
| **Supplementary Figure S6** |  |  |
| Supplementary Figure S6B | *elav<sup>C155</sup>-Gal4/Y;;UAS-PDZD8-V5/ +*<br>*elav<sup>C155</sup>-Gal4/Y;;UAS-PDZD8<sup>SMP\*</sup>-V5/ +* |  |
| Supplementary Figure S6C | *elav<sup>C155</sup>-Gal4/Y;;UAS-PDZD8<sup>SMP\*</sup>-V5/ UAS-Sturkopf-GFP*<br>*elav<sup>C155</sup>-Gal4/Y;;UAS-PDZD8<sup>SMP\*</sup>-V5/ UAS-GFP-Lamp1*<br>*elav<sup>C155</sup>-Gal4/Y;;UAS-PDZD8<sup>SMP\*</sup>-V5/ UAS-GFP-Mito* |  |

## Acknowledgments

We thank the Developmental Studies Hybridoma Bank, the Bloomington Drosophila Stock Center, Dr. Helmut Kramer for sharing p62 antibody, Dr. Pietro De Camilli for providing the hPDZD8-GFP plasmid, and Dr.Gábor Juhász for generously providing antibodies and fly stocks used in preliminary experiments. We thank Dr. Arash Bashirullah, Dr. Raghu Padinjat, Dr. Karla Kaun, Dr. Anup Parchure, Sarah Neuman, Krishna Amin, Kimberly Madhwani, and other members of the O’Connor-Giles lab for thoughtful discussions and comments on the manuscript. This work was supported by a grant from the National Institute of Neurological Disorders and Stroke, National Institutes of Health to K.M.O.G (R01NS078179) and funds from the Brown University Carney Institute for Brain Science

## Author contributions

**Rajan S. Thakur:** Conceptualization; formal analysis; investigation; visualization; writing – original draft; writing – review and editing. **Kate M. O’Connor-Giles:** Conceptualization; formal analysis; funding acquisition; visualization; writing – original draft; project administration; writing – review and editing.

## Declaration of interests

The authors declare no competing interests.

## Inclusion and diversity

Diverse perspectives enhance understanding and accelerate scientific progress. We support scientific practices that broaden participation and inclusion in STEM.

## Notes

### Competing Interest Statement

The authors have declared no competing interest.

### Summary of Updates

References

